# Disrupted Transcriptional Networks by Mutant Atrophin-1 in a Cell Culture Model of Dentatorubral-Pallidoluysian Atrophy

**DOI:** 10.1101/2025.08.08.669318

**Authors:** Oluwademilade Nuga, Masoumeh Pourhadi, Julia P. Rausch, Sokol V. Todi

## Abstract

Dentatorubral-Pallidoluysian Atrophy (DRPLA) is a dominant neurodegenerative disease caused by CAG triplet repeat expansion in *ATN1*, which encodes the transcriptional co-repressor Atrophin-1. DRPLA features motor, cognitive, and epileptic symptoms and shares pathogenic mechanisms with other polyglutamine (polyQ) disorders, including protein misfolding, impaired autophagy, and transcriptional dysregulation. To understand disease mechanisms, we performed RNA-seq on HEK293T cells stably expressing wild-type or polyQ-expanded ATN1. Cells expressing pathogenic ATN1 exhibited a distinct transcriptomic profile, including disruptions in synaptic organization, extracellular matrix remodeling, ion channel expression, and neurotransmission. Several genes tied to neurodevelopmental, neurodegenerative, and oncogenic pathways were fully activated or silenced. Dysregulated pathways also included inflammation, chromatin remodeling, stress responses, and redox imbalance. Heat shock protein expression changes suggested proteotoxic stress and impaired protein quality control, with some findings conserved in a previously reported *Drosophila melanogaster* model of DRPLA. These transcriptomic signatures expand our understanding of molecular events related to degeneration in DRPLA and may lead to the identification of therapeutic targets.

## INTRODUCTION

Dentatorubral-Pallidoluysian Atrophy (DRPLA) is a rare, autosomal dominant neurodegenerative disorder characterized by progressive motor dysfunction, cognitive decline, and epilepsy. Neuropathologically, DRPLA is marked by selective degeneration of the dentatorubral and pallidoluysian systems, neural circuits critically involved in motor coordination and regulation. The disease exhibits geographical clustering, with the highest prevalence reported in Japan and lower incidence observed in European populations (Carroll, Massey, Wardle, & Peall, 2018; Koide et al., 1994; Nagafuchi et al., 1994; Switonska-Kurkowska, Krist, Delimata, & Figiel, 2021).

DRPLA belongs to the polyglutamine (polyQ) family of disorders, a group of inherited, age-dependent neurodegenerative diseases caused by CAG trinucleotide repeat expansions within protein-coding regions. These expansions result in elongated polyQ tracts, leading to protein misfolding and subsequent cellular toxicity. The polyQ disease spectrum includes Huntington’s Disease (HD), Spinal and Bulbar Muscular Atrophy (SBMA), Spinocerebellar Ataxias (SCA) types 1, 2, 3, 6, 7, and 17, in addition to DRPLA. Except for SBMA, which is X-linked, all polyQ disorders follow an autosomal dominant inheritance pattern. A shared clinical hallmark across these disorders is the inverse correlation between CAG repeat length and age of disease onset (Johnson, Tsou, Prifti, Harris, & Todi, 2022; Lieberman, Shakkottai, & Albin, 2019).

In DRPLA, the pathogenic CAG expansion occurs in the *ATN1* gene on chromosome 12, encoding the Atrophin-1 protein. Alleles with more than 48 repeats are typically associated with full disease penetrance. Atrophin-1 is a transcriptional co-repressor that plays essential roles in gene expression regulation. It is part of the evolutionarily conserved Atrophin family of proteins, which includes Atrophin-2 (also known as RERE, or arginine-glutamic acid dipeptide repeat protein). These proteins are implicated in diverse developmental processes across species, including embryonic patterning, neurogenesis, and cardiac development (Shen, Lee, Choe, Zoltewicz, & Peterson, 2007) (Shen & Peterson, 2009; L. Wang & Tsai, 2008; Wood et al., 2000; S. Zhang, Xu, Lee, & Xu, 2002; Zoltewicz, Stewart, Leung, & Peterson, 2004).

Pathogenic, polyQ-expanded Atrophin-1 is thought to exert toxic gain-of-function effects through multiple mechanisms. These include impaired phosphorylation dynamics, which may hinder activation of neuronal survival pathways; defective autophagic processes, which compromise clearance of misfolded proteins; and decreased protein stability, promoting abnormal proteolytic processing and formation of intranuclear inclusions. These molecular events collectively contribute to neuronal dysfunction and cell death (Johnson, Tsou, et al., 2022; Lieberman et al., 2019; Nowak, Kozlowska, Pawlik, & Fiszer, 2023), (Nisoli et al., 2010; Okamura-Oho et al., 2003) (Suzuki, Nakayama, Hashimoto, & Yazawa, 2010).

In addition to its role in DRPLA, mutations in non-expanded regions of *ATN1*, particularly within exon 7, have been associated with *ATN1*-related neurodevelopmental disorder (ATN1-NDD), a distinct condition characterized by intellectual disability, hypotonia, and congenital malformations of the extremities (Whitton, Palmer, & Alkuraya, 1993). Despite growing insights into the molecular pathogenesis of DRPLA and related polyQ disorders, the precise cellular mechanisms remain unclear. At present, there are no disease-modifying therapies available for DRPLA or any other polyQ-associated condition.

Here, we report the generation and transcriptomic characterization of HEK293T cells stably expressing wild-type or pathogenic ATN1. RNA-seq analyses suggest that polyQ-expanded ATN1 disrupts neural gene expression, ion channels, remodeling of the extracellular matrix, stress responses, and redox balance, driving transcriptomic, neurodevelopmental, and proteostasis dysregulation in DRPLA. These and other data may enhance the collective understanding of the biology of disease in DRPLA and pinpoint to potential nodes of intervention.

## MATERIALS AND METHODS

### Cell culture, plasmid, and stable cell preparation

HEK293T cells were obtained from the American Type Culture Collection (ATCC) and maintained in Dulbecco’s Modified Eagle Medium (DMEM, high glucose; ATCC), supplemented with 10% fetal bovine serum (FBS; Gibco) and 1% Penicillin-Streptomycin solution (Gibco). Cells were cultured in a humidified incubator at 37°C with 5% CO₂. Cultures were passaged at 70–80% confluency using 0.25% trypsin-EDTA (Gibco).

Human, wild-type ATN1 cDNA encoding 7 glutamines (Q7-WT) and mutant ATN1 cDNA encoding 88 glutamines (Q88-DRPLA) were synthesized and cloned into pINTO mammalian expression vector (figure 1A) by GenScript (www.genscript.com). A vector lacking any exogenous insert was used as a negative control. The cDNA sequences were derived from Ensembl (<u>ENST00000396684.3</u>) with a CAGCAA repeat sequence replacing the pure CAG tract. This modification was implemented to avoid repeat-associated non-ATG (RAN) translation and reduce the risk of RNA-mediated toxicity typically associated with uninterrupted CAG repeats. (Johnson, Prifti, et al., 2022; L. B. Li, Xu, & Bonini, 2007; L. B. Li, Yu, Teng, & Bonini, 2008; Pearson, 2011; Sobczak, de Mezer, Michlewski, Krol, & Krzyzosiak, 2003; Sobczak & Krzyzosiak, 2004a, 2004b, 2005; Zu, Pattamatta, & Ranum, 2018). Each construct was sequence-verified before utilization for its intended purposes.

**Figure 1.**
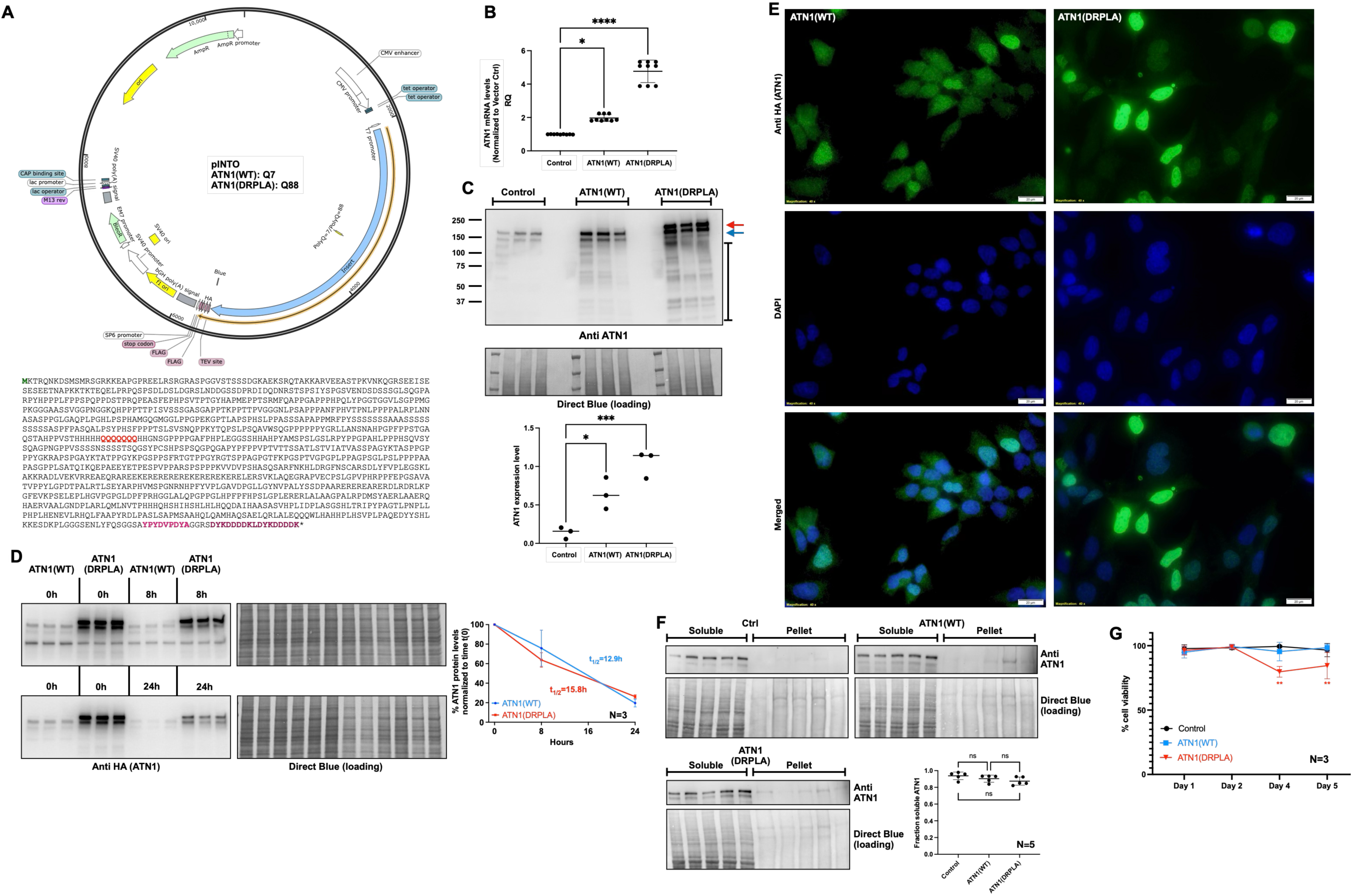
Generation of HEK-203T cells stably over-expressing ATN1. (A) Schematic and features of the pINTO plasmid and the amino acid sequence of the encoded ATN1 protein. Green font, starting methionine; red font, the polyQ region that is represented by 88Q in the pathogenic version of the protein used in this study; shades of purple, the HA (first) and FLAG (second) tags. (B) qPCR of the expression levels of endogenous and stably expressed ATN1 constructs. Shown are means -/+ SD. Statistics, Kruskal-Wallis. *, P<0.05, ****, P<0.0001. N=3 independent repeats. (C) WB HEK293T cells expressing the noted constructs. Blue arrow, ATN1(WT) which migrates similarly to endogenous ATN1. Red arrow, ATN1(DRPLA). Bracketed line, likely proteolytic fragments of ATN1. Statistics in quantification, one-way ANOVA. Shown are means -/+ SD. *, P<0.05, ***, P<0.001. N=3 independent repeats. (D) WB from CHX-based pulse/chase assays of the noted HEK293T cells. Quantifications are means -/+ SD. Non-statistically significant by two-way ANOVA. N=3 independent repeats. (E) Epi-fluorescence images of the indicated cells, imaged with an Olympus BX53 microscope at 40X magnification. Scale bar on the right side, 20 µm. (F) Soluble/pellet separation of ATN1 protein from the indicated cells. Shown in quantifications are means -/+ SD, where ATN1 signal, which was normalized to its respective Direct Blue loading, was mathematically adjusted to reflect the fraction of total sample it represents. Statistics are from Brown-Forsythe and Welch ANOVA. ns, not significant. N=5 independent repeats. (G) MTT cell viability assays of the indicated cells. Means -/+ SD. Statistics, two-way ANOVA. **, P<0.01. N=3 independent repeats.

HEK293T cells were seeded at a density of 2.5 × 10⁵ cells per well in 12-well plates and cultured in antibiotic-free medium. Cells were transfected with 0.5 µg of plasmid DNA per well using Lipofectamine™ LTX with PLUS™ Reagent (Thermo Fisher Scientific), following the manufacturer’s instructions. For each transfection, the DNA was first combined with PLUS™ Reagent, incubated, and then mixed with Lipofectamine™ LTX to form DNA-lipid complexes, which were added directly to the cells. An empty vector was used in parallel as a negative control. Cells were selected with Gibco Zeocin Selection Reagent (Thermo Fisher Scientific) at a concentration of 400 µg/mL 48 hours post-transfection for 10 days. Established stable cell lines were maintained in medium containing 100 µg/mL zeocin to preserve selective pressure (Lanza, Kim, & Alper, 2013). The Zeocin concentration was determined based on a kill curve performed to identify the dose required for complete cell death in non-transfected HEK293T cells. To establish stable cell lines, transfected Individual resistant colonies were manually picked, expanded, and screened for transgene expression by Western blotting.

### Cell viability

Stable cell lines expressing empty vector control, ATN1(Q7-WT), or ATN1(Q88-DRPLA) were seeded in 96-well plates at a density of 7×10^3^ cells/well. Cell viability was assessed daily using the MTT assay. At each time point, 10 µL of MTT solution (5 mg/mL in PBS, Merck) was added to each well, and cells were incubated for 3–4 hours at 37°C. After incubation, 100 µL of DMSO was added to solubilize the formazan crystals, and absorbance was measured at 570 nm using Tecan Infinite M Plex microplate reader (Pearson, 2011). Viability was calculated as relative absorbance compared to the control group on the same day.

### qRT-PCR

Total RNA was extracted from stable cell lines using the PureLink™ RNA Mini Kit (Thermo Fisher Scientific) and genomic DNA was eliminated by treatment with TURBO DNase (Thermo Fisher Scientific). Reverse transcription was carried out using the high-capacity cDNA reverse transcription kit (ABI). mRNA levels were quantified using the StepOne Real-Time PCR system with Fast SYBR Green Master Mix (ABI). Primers used were:

ATN1 F: 5′ - AGTGATGGCAAAGCTGAGAAGT 3′;

ATN1R: 5′ - CTTCCAGGGCTGTAGATACTGG- 3

GAPDH-F: 5′-CTCAGACACCATGGGGAAGGT-3′;

GAPDH-R: 5′-GTGGTGCAGGAGGCATTGCTGA-

### Cycloheximide assay

Cells were seeded at a density of 3 × 10⁵ cells per well in 12-well plates and incubated overnight to allow attachment. The next day, cycloheximide (CHX, GoldBio, 1203.030519A) with the final concentration of 50 µM was added to the cell culture medium to inhibit new protein synthesis. Cells were collected at intervals 0-, 8-, and 24-hours post-treatment for protein stability analysis. Protein half-life (t½) was calculated from densitometric values obtained from Western blots.

### Rotenone-induced cytotoxicity

Stable cell lines were treated with increasing concentrations of rotenone (0, 1, 10, 20, 50, 100, 250, and 500 nM, Chem Cruz) to assess rotenone-induced toxicity. Cells were seeded in 96-well plates at a density of 7×10^3^ cells/well in triplicate and allowed to adhere overnight. Rotenone treatments were applied for 24 and 48 hours. Following treatment, cell viability was assessed via MTT assay as described above. Formazan crystals were solubilized with DMSO, and absorbance was measured at 570 nm. Data were normalized to the untreated control (0 nM).

### RNA Extraction, sequencing, and bioinformatics

Stable cell lines at passage 13 (five passage post-selection), expressing empty control vector, ATN1(WT) or ATN1(DRPLA) were used for RNA extraction and sequencing, with 5 biological replicates per group. Cells were cultured under standard conditions and total RNA was extracted using the PureLink™ RNA Mini Kit (Thermo Fisher Scientific) according to the manufacturer’s instructions and genomic DNA was eliminated by treatment with TURBO DNase (Thermo Fisher Scientific). RNA quantity and purity were assessed by NanoDrop, and integrity was checked using a Bioanalyzer (Agilent Technologies; supplemental figure 1). RNA samples were submitted to Azenta Life Sciences for RNA sequencing and library preparation.

Samples underwent standard RNA sequencing with rRNA depletion, library preparation, and sequencing on the Illumina platform with paired-end 150 bp reads (2 × 150 bp). A sequencing depth of approximately 30 million reads per sample was achieved. Sequencing adaptors were trimmed, and unique molecular identifier (UMI)-based de-duplication was carried out simultaneously using fastp v0.23.1. Trimmed and de-duplicated reads were aligned to the Homo sapiens GRCh38 reference genome (ENSEMBL release) using the STAR aligner v2.5.2b. Unique gene-level counts were obtained using the featureCounts function from the Subread package v1.5.2. Read quality was initially assessed using FastQC, and downstream quantification and statistical analysis were conducted in R.

Differential expression analysis was performed with DESeq2(Love, Huber, & Anders, 2014), applying a Wald test to compute p-values and log2 fold changes. Genes were considered differentially expressed if they met the criteria of false discovery rate (FDR) < 0.05 and absolute log2 fold change > 1. Gene Ontology (GO) analysis of differentially expressed genes was conducted using the ‘clusterProfiler’ package (v4.6.2) in R. For functional enrichment and gene set enrichment analysis (GSEA), the Enrichr webtool was used (Chen et al., 2013). Analyses included the enrichment of transcriptional regulators and biological pathways to contextualize the observed gene expression changes.

### Antibodies

Anti-ATN1 mouse monoclonal (Santa Cruz, Cat. #sc-517594), 1:200 for WB; anti-HA, rabbit monoclonal (Cell Signaling, Cat. #372), 1:1000 for WB and IFC; anti AR, rabbit monoclonal (Cell Signaling, Cat #5153), 1:500 for WB, Anti ATXN2, rabbit monoclonal (Cell Signaling, Cat #35121), 1:1000 for WB; anti ATXN3 (MJD), rabbit polyclonal (MJD), rabbit polyclonal (Paulson et al., 1997) (REF), 1:5000 for WB; anti-ATXN7, rabbit polyclonal (Mohan et al., 2014), 1:1000 for WB; anti MAGE-A3, rabbit polyclonal (Cell Signaling, #25800), 1:1000 for WB goat anti-rabbit, Alexa Fluor™ Plus 488, (ThermoFisher Catalog #: A-11034), 1:1000 for microscopy; goat anti-rabbit peroxidase-conjugated secondary and goat anti-mouse HRP secondary (Jackson Laboratories, Cat.#115–035–144 and 115–035–146; 1:5000).

### Protein extraction and Western blotting

Cells were washed with phosphate-buffered saline (PBS; Gibco) and lysed with boiling lysis buffer (61.2% 2x Laemmli Buffer, 30.1% MilliQ H2O, 8.2% Dithiothreitol (DTT), 0.01g Bromophenol Blue Dye), followed by boiling for 7 minutes and centrifuged at 13,300 x g for 4 minutes at room temperature. The samples were loaded into pre-cast 4–20% Tris/Glycine gels (Bio-Rad) to separate proteins by electrophoresis and then transferred onto PDVF membranes (Bio-Rad). The membranes were blocked for 30 minutes using EveryBlot Blocking Buffer (Bio-Rad) and probed with primary antibody overnight at 4°C. An HRP-conjugated secondary antibody was introduced for 1 hour at room temperature. Imaging was performed using the ChemiDoc Imaging System (Bio-Rad). The membranes were stained with Direct Blue 71 (40% ethanol, 10% acetic acid, 50% MilliQ H2O, .0008% Direct Blue 71 (Sigma Aldrich)) for 10 minutes, followed by rinsing with 40% ethanol, 10% acetic acid, and 50% MilliQ H2O. The membranes were air dried and imaged using ChemiDoc. Bands from Western blots and total protein (Direct Blue) staining were quantified using ImageLab software (Bio-Rad).

### Soluble/pellet preparation

For the separation of cellular proteins into soluble/pellet fractions, adherent cells were washed with ice cold phosphate-buffered saline (PBS; Gibco) with PMSF and harvested using either NETN (50mM Tris, pH 7.5, 150 mM NaCl, 0.5% Nonidet P-40) or RIPA (50 mM Tris, 150 mM NaCl, 0.1% SDS, 0.5% deoxycholic acid, 1% Nonidet P-40, pH 7.4) lysis buffer supplemented with PMSF, protease inhibitor cocktail V (Calbiochem) and phosphatase inhibitor III (Calbiochem). Lysates were then sonicated (50% power for 15 seconds on ice). Samples were then centrifuged at 20,000 x g at 4°C for 30 minutes and the supernatant (soluble fraction) was transferred into a new microfuge tube on ice. The pellet was rinsed again, vortexed lightly with RIPA buffer and centrifuged at 20,000 x g at 4°C. Supernatant was aspirated completely leaving pelleted fraction. Pellet was then resuspended in 50 uL Laemmli buffer (12.5% 1M Tris, 20% Glycerol, 4% SDS, pH 7.5) and sonicated again at 50% AMP for 30 seconds. The resulting samples were then centrifuged for 2 minutes at 1000 x g and both soluble and pellet portions were supplemented with 6% SDS. Samples were then boiled 10 min, spun down briefly and loaded onto SDS-PAGE gels.

### Cell imaging

HEK293T cells were seeded in 12-well plates on poly-D-lysine-coated coverslips and cultured overnight to allow attachment. Cells were then fixed with 4% paraformaldehyde in PBS for 15 minutes, permeabilized with 0.1% Triton X-100, and blocked in EveryBlot Blocking Buffer (Bio-Rad). Primary antibody was applied overnight at 4°C, followed by Alexa Fluor™ Plus 488-conjugated goat anti-rabbit IgG secondary antibody for 1 hour at room temperature. Coverslips were mounted using ProLong™ Diamond Antifade Mountant with DAPI (Invitrogen). Cells were imaged at 40X magnification using an Olympus BX53 microscope.

### Statistics

Statistical analyses were conducted through GraphPad Prism version 9 and 10. Other unique analyses are noted in the respective figure legends throughout the paper.

## RESULTS

### Stable expression of wild-type and pathogenic ATN1 in HEK293T cells

Towards understanding the biology of disease of DRPLA and finding potential therapeutic routes for it, we generated HEK293T cells that stably express an empty vector control, wild-type ATN1 or pathogenic ATN1. We selected HEK293T cells due to their ease of maintenance, manipulation, and future usability in large screens of pharmaceutical compounds and other types of investigations (Pulix, Lukashchuk, Smith, & Dickson, 2021; Stepanenko & Dmitrenko, 2015; Tan, Chin, Lim, & Ng, 2021; Thomas & Smart, 2005). The sequence of the resulting protein is shown in figure 1A. We selected a wild-type sequence of 7Q and a pathogenic one of 88Q, within patient range (Johnson, Tsou, et al., 2022). As noted in figure 1A, the ATN1 constructs include HA and FLAG epitope tags at their C-termini.

We validated transgene expression through quantitative PCR (qPCR), Western blotting (WB) and microscopy. Results were consistent across independent replicates (figure 1) and multiple passage numbers (data not shown), confirming the stability of the expression system. Based on qRT-PCR, exogenous ATN1 is expressed at higher levels than the endogenous gene, with the pathogenic version (ATN1(DRPLA)) being highest (figure 1B). This difference in expression was confirmed by WBs, where pathogenic ATN1 was again at higher levels than overexpressed ATN1(WT) or its endogenous counterpart (figure 1C). In WBs, the full-length ATN1 bands appear in the expected molecular weight range of 150–200 kDa, with the mutant version migrating more slowly than the wild-type due to the expanded polyQ tract. Besides the primary band, indicated by arrows in figure 1C, we observed additional ATN1-positive signal, likely consistent with proteolytic fragments, which have been reported before for this protein (Ellerby et al., 1999; Nucifora et al., 2003; Suzuki et al., 2010). Because of the higher levels of ATN1(DRPLA) compared to its (WT), overexpressed counterpart, we investigated if the turnover rates of these proteins were different in these cells. We used CHX-based assays to stop overall protein translation for 0, 8 or 24h, and found that there was no discernible difference in their turnover (figure 1D).

Next, we examined the localization of the stably expressed proteins through epi-fluorescence microscopy. In ATN1(WT)-expressing cells, the protein shows a diffuse distribution in both the cytoplasm and nucleus; some cells show nuclear accumulation (figure 1E and supplemental figure 2A). ATN1(DRPLA)-expressing cells show higher overall levels and with more frequent accumulation in nuclei (figure 1E and supplemental figure 2A). We did not observe frequent indications of aggregation in HEK293T cells expressing ATN1(DRPLA): examples in figure 1E do not show indications of aggregations; examples in supplemental figure 2A show instances of aggregation, but these were not common.

Because mutant ATN1 aggregates in DRPLA patient tissue and in animal models of this disease (Carroll et al., 2018; Koide et al., 1994; Nagafuchi et al., 1994; Switonska-Kurkowska et al., 2021) (Prifti et al., 2025) (Napoletano et al., 2011; Nisoli et al., 2010; Zoltewicz et al., 2004), we conducted biochemical, soluble/pellet centrifugation of stably expressing cells. As summarized in figure 1F, ATN1 signal was detected in the soluble fraction for both constructs, but there were also bands in the pellet fraction (figure 1F). We observed similar outcomes whether using the more stringent RIPA lysis buffer (figure 1F) or the milder, NETN lysis buffer (supplemental figure 2B). Based on the quantification of these data, there were no significant differences in the fraction of soluble versus pellet signal among the three groups in either buffer (figure 1F, supplemental figure 2B).

Finally, we tested if stable over-expression of ATN1(WT) or (DRPLA) is toxic to HEK293T cells. We conducted MTT assays over a 5-day period. All groups maintained high viability (>96%) during days 1 and 2. However, by day 4 and day 5, we observed reduced viability in cells expressing ATN1(DRPLA) compared to both wild-type and control cells (figure 1G). Altogether, these data indicate that both wild-type and pathogenic ATN1 proteins are stably expressed in HEK293T cells, that the proteins are largely soluble in this model, and that expression of polyQ-expanded ATN1 may induce cytotoxic stress over time.

### Global transcriptomic changes

We profiled the transcriptome of HEK293T cells expressing control, ATN1(WT), or ATN1(DRPLA) by performing bulk paired end RNA-seq. Outlined in figure 2A is the analysis pipeline. The principal component analysis (PCA) showed clear separation of DRPLA samples from both control and ATN1(WT), indicating a distinct transcriptomic profile (figure 2B). We then performed a comparative differential expression analysis to identify genes that were significantly changed in between conditions with an absolute log2FC cutoff of 1. Focusing on differentially expressed genes (DEGs) in ATN1(DRPLA) versus ATN1(WT), we identified 1,791 genes that were differentially expressed, with 479 genes uniquely altered in ATN1(DRPLA) compared to ATN1(WT), while other DEGs were similarly deregulated between ATN1(WT) and control or ATN1(DRPLA) and control (figure 2C; supplemental tables 1 and 2).

**Figure 2.**
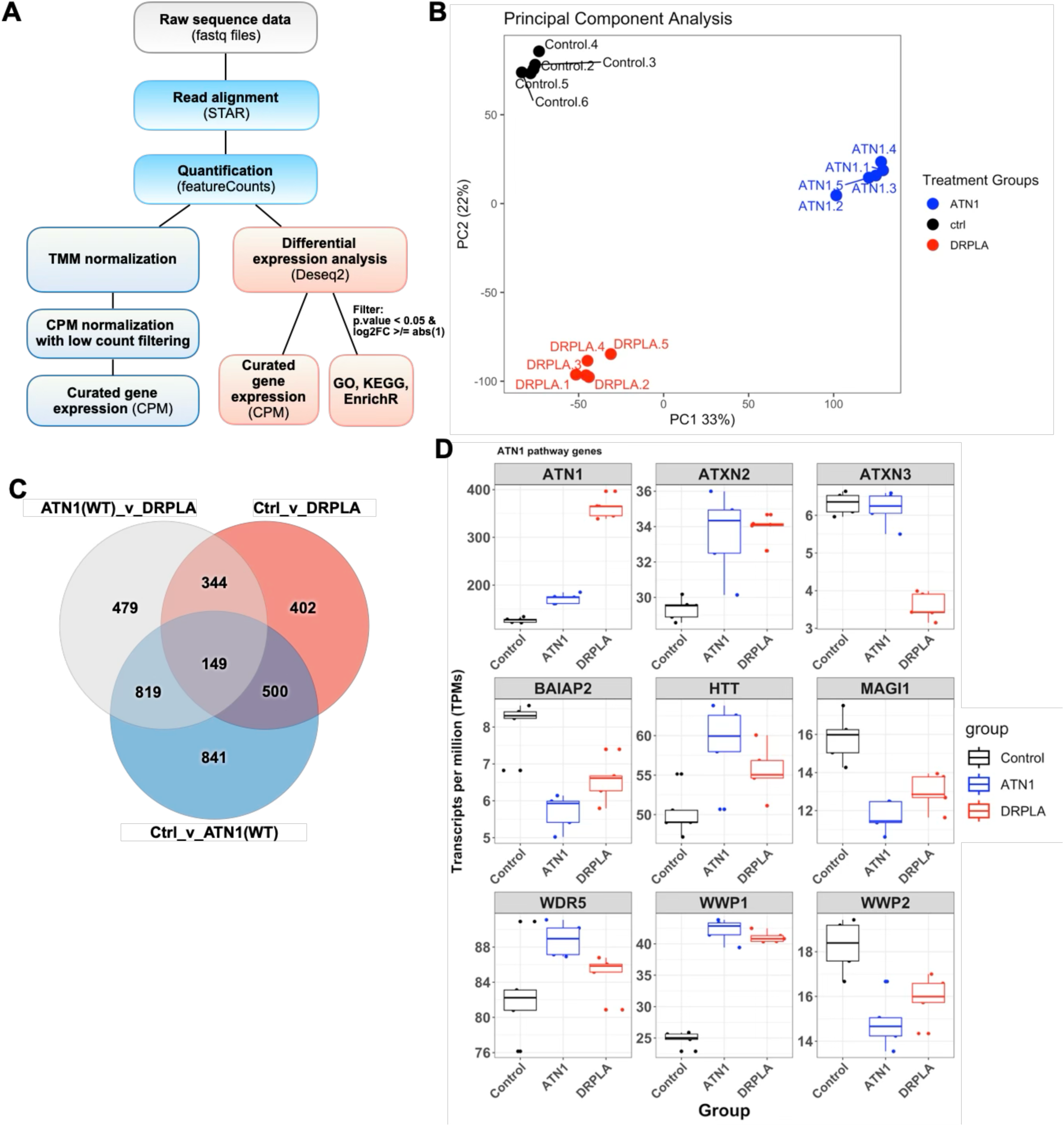
Transcriptomic profiling reveals distinct gene expression patterns among the stable HEK293T cells. (A) Schematic of the RNA-seq analysis pipeline comparing HEK293T cells stably expressing control vector, ATN1(WT), or ATN1(DRPLA), N=5 independent replicates. (B) Principal component analysis (PCA) performed using transcript per million (TPM) values demonstrates distinct separation of ATN1(DRPLA) samples from both ATN1(WT) and control groups, indicating a unique transcriptional signature driven by the polyQ expansion. (C) Venn diagram summarizing the number of differentially expressed genes (DEGs; absolute log2FC ≥ 1) across comparisons including ‘control versus ATN1(WT)’, ‘control *versus* ATN1(DRPLA)’ and ‘ATN1(WT) *versus* ATN1(DRPLA)’. A total of 1,791 DEGs were identified in ATN1(DRPLA) versus ATN1(WT), including 479 genes that did not overlap with other comparisons. (D) Boxplot of mRNA expression changes (TPM) of selected genes involved in ATN1-associated regulatory pathways.

Given the role of ATN1 as a transcriptional co-repressor with endogenous function in HEK-293T cells, we assessed whether components of the ATN1-associated regulatory network responded to increases in wild-type (mRNA fold change < 2) or pathogenic ATN1 expression (mRNA fold change < 6) (figure 1B). Relative to the control, Ataxin-2 (ATXN2, which causes SCA2), WRD5, Huntingtin (HTT, which causes HD), and WWP1 were upregulated in both ATN1(WT), and ATN1(DRPLA) cell lines (figure 2D), suggesting that these genes are responsive to elevated ATN1 levels regardless of polyQ expansion. In contrast, BAIAP2, MAGI1, and WWP2 were consistently downregulated in both conditions (figure 2D), indicating potential transcriptional repression or loss of scaffolding regulation linked to ATN1 overexpression.

Intrigued by the changes in mRNA levels of the polyQ disease protein-encoding genes in figure 2D, we analyzed the impact of the exogenous, WT or DRPLA version of ATN1 on all polyQ disease-encoding genes in these samples (supplemental figure 3). We noted that nearly every polyQ disease gene was impacted in one direction or another by ATN1(WT) or (DRPLA) in these samples. Levels of AR (which causes SBMA) and ATXN2 were increased in the presence of pathogenic ATN1; we confirmed these findings with WB (supplemental figure 4). ATXN3 (which causes SCA3) mRNA expression was unchanged in ATN1(WT) but significantly reduced in ATN1(DRPLA), suggesting that the expanded polyQ tract of ATN1 disrupts ATXN3 expression (figure 2D). However, when probing for the protein levels of ATXN3, we observed its increased overall protein levels in ATN1(DRPLA) cells and a marked change in the intensity of the higher molecular weight species of this protein (supplemental figure 4), which are concomitant with its posttranslational modification by ubiquitination and phosphorylation (Johnson, Tsou, et al., 2022). The incongruent results for ATXN3 mRNA *versus* protein levels highlights the need to conduct future, in-depth proteomics work with the stable HEK293T cells to understand the coordination of overall cellular processes altered by the presence of normal or pathogenic ATN1. ATXN7 (which causes SCA7) levels were decreased at the mRNA and protein level in the presence of ATN1(DRPLA); the results were incongruent regarding the impact of ATN1(WT) on ATXN7 levels, which were increased at the mRNA level did not reach significant difference at the protein level (supplemental figure 2). Insofar as the other polyQ diseases are concerned, ATXN1 (which causes SCA1) mRNA levels were higher in both ATN1(WT) and ATN1(DRPLA) groups; CACNA1A (whose C-terminal mutation causes SCA6) was not markedly impacted (but, please note that the reads of CACNA1A were very low and did not pass quality control; they are shown here for the sake of completeness); TBP (which causes SCA17) levels were increased in both ATN1(WT) and ATN1(DRPLA) groups compared to the control. Based on these results, we conclude that ATN1 over-expression leads to polyQ-dependent and -independent alterations in the transcriptome of HEK293T cells and may have particular importance to the regulation of polyQ disease protein-encoding genes.

### Dysregulated pathways and biological processes

To understand how polyQ-expanded ATN1 alters cellular programs, we analyzed gene expression changes associated with biological processes (figure 3). Relative to ATN1(WT), expression of pathogenic ATN1 resulted in downregulation of genes involved in synapse organization, axonogenesis, and cell adhesion, all key pathways supporting neural connectivity. In contrast, transcripts related to integrin-mediated adhesion and extracellular matrix (ECM) remodeling were increased, indicating a shift away from neural-specific signaling and toward structural reorganization (figure 3A). Notably, components of synaptic transmission appeared in both upregulated and downregulated gene sets in this comparison, consistent with broader disruption of synaptic regulatory balance.

**Figure 3.**
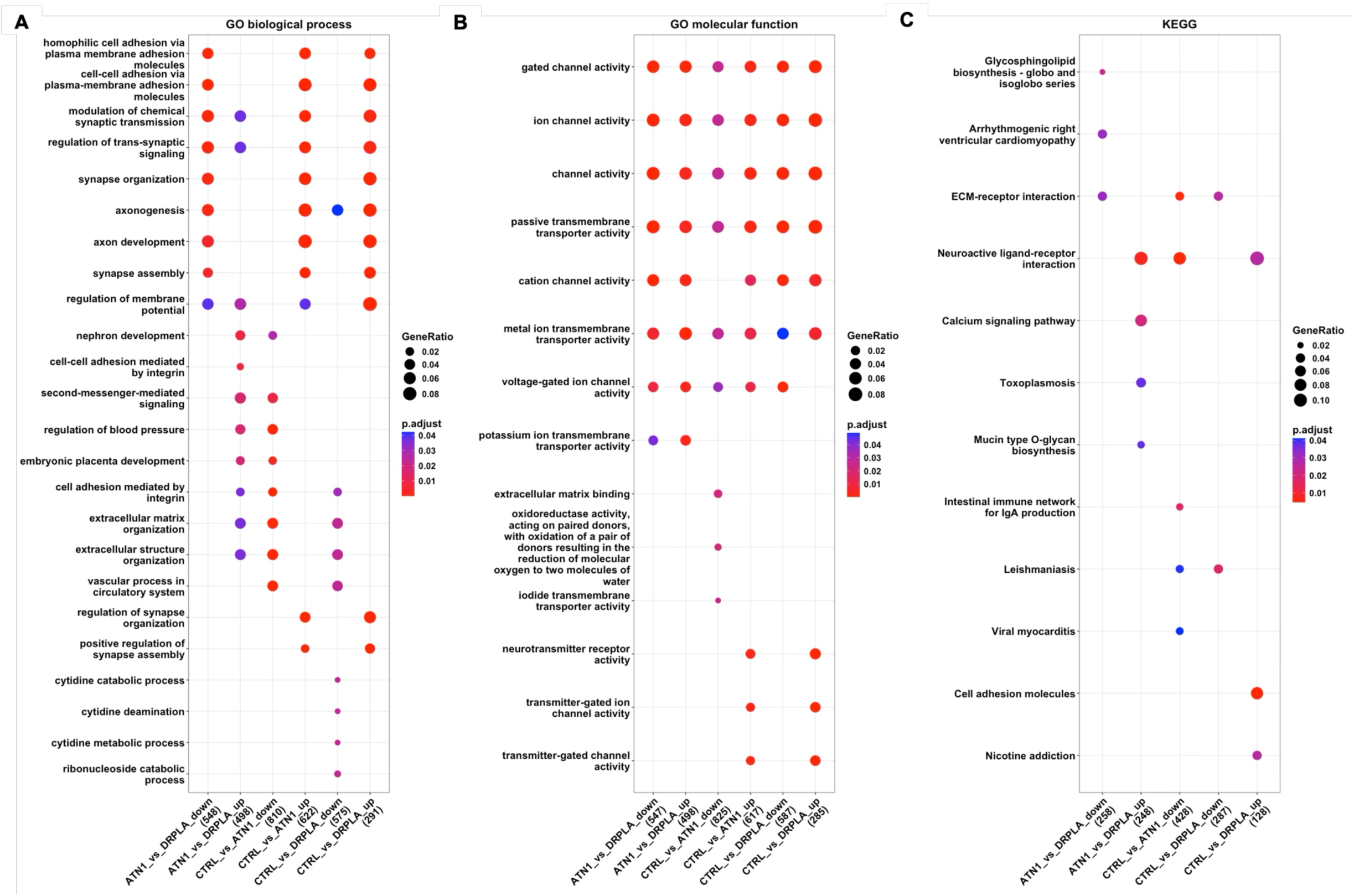
Functional enrichment analyses of differentially expressed genes across ATN1 expression conditions. Gene ontology (GO) and KEGG pathway enrichment analyses were performed on up- and downregulated differentially expressed genes identified in the following pairwise comparisons: control *versus* ATN1(WT), control *versus*. ATN1(DRPLA), and ATN1(WT) *versus* ATN1(DRPLA). (A) GO Biological Process terms. (B) GO Molecular Function terms. (C) KEGG pathway analysis.

The ATN1(WT) *versus* vector control comparison revealed the opposite trend: genes linked to axon formation and synapse assembly were upregulated, while ECM- and integrin-associated adhesion genes were suppressed (figure 3A). These results support a potential role for wild-type ATN1 in maintaining neural-like transcriptional programs, even in non-neuronal cells such as HEK293T. In the ATN1(DRPLA) versus control comparison, both neural and non-neural pathways were affected. Genes promoting axonal growth, synaptic organization, and membrane potential regulation were increased, whereas those involved in ECM maintenance, structural adhesion, and nucleotide metabolism were decreased (figure 3A).

Analysis of molecular function categories pointed to broad dysregulation of ion channel activity and neurotransmission in the DRPLA model (figure 3B). Compared to ATN1(WT), ATN1(DRPLA) induced upregulation of multiple voltage-gated ion channels, including calcium (CACNA1G, CACNA1I), sodium (SCN2A, SCN3A), and potassium (KCNQ4, KCNC4, KCNV1) channels, as well as glutamate and GABA receptor subunits (GRIN1, GABRE, GABRA2, GRIK3; supplemental tables 1, 2). Expression of mechanosensitive and calcium-release channels (PIEZO1, TRPV4, RYR2) was also elevated, suggesting increased sensitivity to cellular stress and altered calcium handling (supplemental tables 1, 2). At the same time, transcripts encoding regulators of ion conductance and neurotransmission were downregulated, including ABCC8 and ABCC9 (involved in potassium channel modulation), TRPM5 (a calcium-activated channel), and glutamate receptors (GRIK5, GRIA4), along with several GABA and cholinergic receptor subunits (GLRA1, CHRND, CHRNB2, CHRNA9; supplemental tables 1, 2). Downregulation of key potassium channel components, including KCNA1, KCNA2, and KCNMA1, may impair repolarization capacity, further increasing excitability in a dysregulated context.

We extended this analysis using KEGG pathway annotation (figure 3C). Genes involved in glycosphingolipid biosynthesis and ECM-receptor interactions were consistently upregulated in ATN1(DRPLA) relative to ATN1(WT), and ECM-related downregulation was also evident when comparing ATN1(DRPLA) to vector control. Neuroactive ligand–receptor signaling pathways showed opposing regulation: downregulated in ATN1(WT) versus control, but upregulated in ATN1(DRPLA), suggesting a reversal of normal signaling dynamics. Additionally, expression of cell adhesion molecules increased in ATN1(DRPLA) relative to control (figure 3C).

Collectively, these findings indicate that polyQ expansion in ATN1 perturbs the balance between structural signaling and neural communication. Dysregulation of glycosphingolipid metabolism, excitatory/inhibitory imbalance, ion channel dysfunction, and cytoskeletal remodeling likely contribute to the disease-relevant cellular phenotypes observed in this model.

### Binary gene expression shifts in ATN1(WT) versus (DRPLA)

We next examined genes that were fully turned “on” or “off” in ATN1(WT) or ATN1(DRPLA), since binary expression changes can signal critical regulatory shifts triggering or silencing entire pathways, acting as molecular switches with disproportionate influence on cellular state. For these investigations, we first focused on comparisons where ATN1(DRPLA) results differed from both Ctrl and ATN1(WT), which were similar to each other.

We recovered a set of “turned on/off” genes from our transcriptomic screen that are implicated in diverse neurological and oncological conditions (Figure 4). A broader inspection of these data reveals that the pattern of binary expression changes in genes either fully activated or silenced varies across the three experimental groups (Figure 4A). Specifically, some gene clusters show expression similarities between ATN1(WT) and ATN1(DRPLA) relative to control cells, pointing to transcriptional programs that are dependent on ATN1 overexpression but not necessarily affected by the polyQ expansion itself. Several genes within this group have documented functions in synaptic signaling, neurodevelopment, and inflammatory processes. These include ASTN1 (neuronal migration), CHRM3 (cholinergic signaling), DOK6 and DMRTA2 (axon development), ZNF804A and IGSF11 (synaptic plasticity), NPY1R (neuropeptide signaling), PTPRZ1 (glial regulation), MYO7A and MYD88 (neuroimmune signaling), and DNAJC5G (chaperone activity).

**Figure 4.**
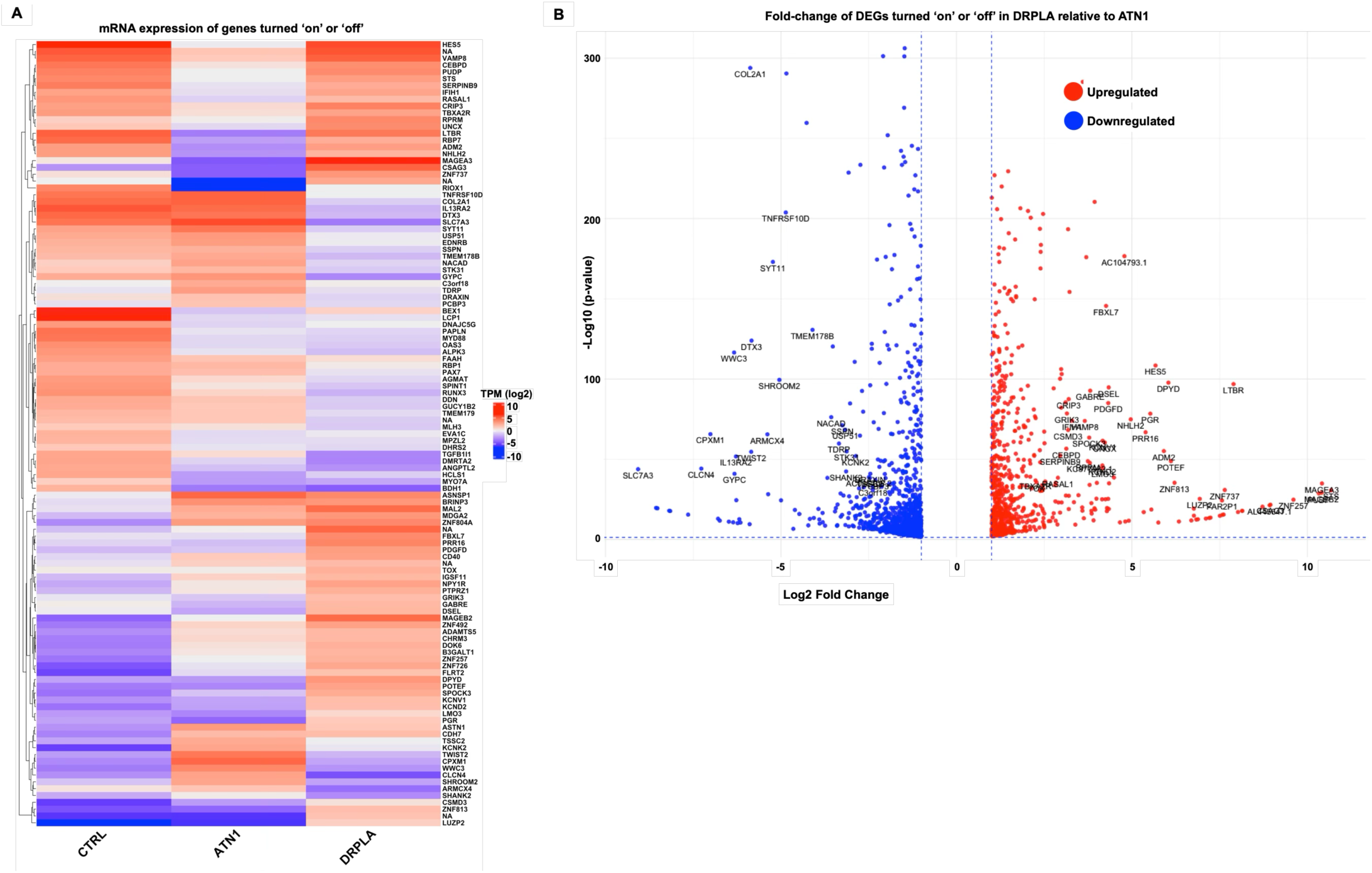
Gene-level transcriptional activation and silencing in polyQ-expanded ATN1. (A) Heat-map of mRNA expression of genes with <25 TPMs in any one condition and a corresponding expression of 100 TPMs in another, across vector control, ATN1(WT), and ATN1(DRPLA) samples. Unsupervised hierarchical clustering separates ATN1(DRPLA) from WT and control, with blocks of genes showing coordinated activation- (red) or silenced-regulation (blue) in the DRPLA condition. (B) Volcano plot comparing ATN1(DRPLA) to ATN1(WT). The X-axis depicts log2 fold-change (log2FC) in ATN1(DRPLA) relative to ATN1(WT), and the Y-axis shows –log10 adjusted p-value. Genes meeting significance (|log2FC| ≥ 1, FDR < 0.05) are colored: red, transcriptionally activated (“turned on”); blue, transcriptionally silenced (“turned off”). Genes that are activated or silence are text annotated in plot. Together, the heat-map and volcano plot highlight the broad transcriptional reprogramming and discrete gene-level shifts driven by polyQ expansion of ATN1.

In contrast, another subset of genes shows expression patterns that are similar between ATN1(WT) and control samples but are altered in ATN1(DRPLA), indicating transcriptional programs that remain stable in the presence of wild-type ATN1 but are perturbed by the polyQ expansion. These include genes involved in chromatin organization, stress responses, synaptic transmission, and signaling pathways. Examples include SYT11 and GRIK3 (involved in vesicle trafficking and glutamatergic signaling), KCNV1 and KCND2 (potassium channels), PAX7, RUNX3, and LMO3 (developmental transcription factors), USP51 and MLH3 (DNA damage repair), PDGFD and EDNRB (growth factor signaling), FAAH and SPINT1 (enzymes involved in lipid metabolism and protease inhibition) and DRAXIN (axon guidance). KCND2, CSMD3, GRIK3, KCNV1, and DRAXIN have prior links to autism spectrum disorder (ASD) and neurodevelopmental delay. Others, including PDGFD and TOX, are associated with glioblastoma progression and tumor immune evasion. In addition, cancer/testis antigens MAGEA3, MAGEB2, and CSAG3, which are aberrantly expressed in brain tumors/melanomas and considered immunotherapy targets, were part of this binary set. Notably, MAGEA3 exhibited concordant changes in both mRNA and protein levels (supplemental figure 4), highlighting its potential translational relevance.

Conversely, there are also genes whose levels are impacted by the over-expression of ATN1(WT), compared to Ctrl samples, but their expression levels are reverted towards normalcy by DRPLA mutations. These may represent transcriptional programs that are actively induced by wild-type ATN1 and then suppressed or reverted by the polyQ-expanded version, suggesting a loss-of-function effect. This group includes transcriptional regulators such as HES5, NHLH2, UNCX, TWIST2, and TBXA2R; WWC3, a Hippo pathway modulator; immune and inflammation-related genes like CEBPD, IFIH1 (linked to interferon-mediated neurodegeneration), LTBR, and SERPINB9; chromatin and DNA damage response components including RPRM and RIOX1; and structural or synaptic genes such as SHROOM2, SHANK2, and KCNK2 (TREK-1), a potassium channel with neuroprotective properties.

Focusing on the loss- or gain-of-function in ATN1(DRPLA) versus ATN1(WT), we observed repressed genes across categories that are relevant to neurodevelopment and tumor biology. DRAXIN, a guidance cue implicated in ASD, was significantly downregulated, pointing to potential disruption in neuronal connectivity. Suppression of TNFRSF10D, an anti-apoptotic decoy receptor, and DTX3, an E3 ligase involved in Notch signaling, suggests dysregulation of apoptotic and differentiation pathways. Additional downregulated genes include SYT11 and SSPN (synaptic vesicle trafficking and cytoskeletal anchoring), COL2A1 (extracellular matrix), and USP51, which has been recently linked to Non-Syndromic X-linked Intellectual Disability 99 (figure 4B).

Together, the binary changes in these and other genes highlights the possibility of coordinated disruption of pathways critical for neuronal connectivity, synaptic stability, apoptosis regulation, and neurodevelopmental integrity in DRPLA (figure 4). Furthermore, there is evidence of neurodevelopmental issues in polyQ diseases, including HD and SCA1 (Blum, Schwendeman, & Shaham, 2013; Edamakanti, Do, Didonna, Martina, & Opal, 2018; Humbert, 2010; Jain et al., 2023; Lieberman et al., 2019; Ratie & Humbert, 2024; Switonska-Kurkowska et al., 2021), pointing to the need for developmental research in the broader context of polyQ disorders.

### Transcriptional network dysregulation in DRPLA

To assess the disease relevance of our findings, we next compared our dataset to publicly available transcriptional profiles using ENRICHR. Hexagonal canvas visualization revealed enrichment of gene sets associated with activation or overexpression of transcriptional regulators such as NFKB1, USF1, TP63, and IRF1 (figure 5A). Additionally, signatures corresponding to DNMT1 inhibition, MYC knockdown, and NCOA1 knockout were upregulated and showed concordant expression with differentially expressed genes (DEGs) in ATN1(DRPLA) versus ATN1(WT) cells (figure 5A, B). These patterns suggest coordinated activation of pathways involved in inflammation, chromatin remodeling, cell cycle regulation, and transcriptional coactivation.

**Figure 5.**
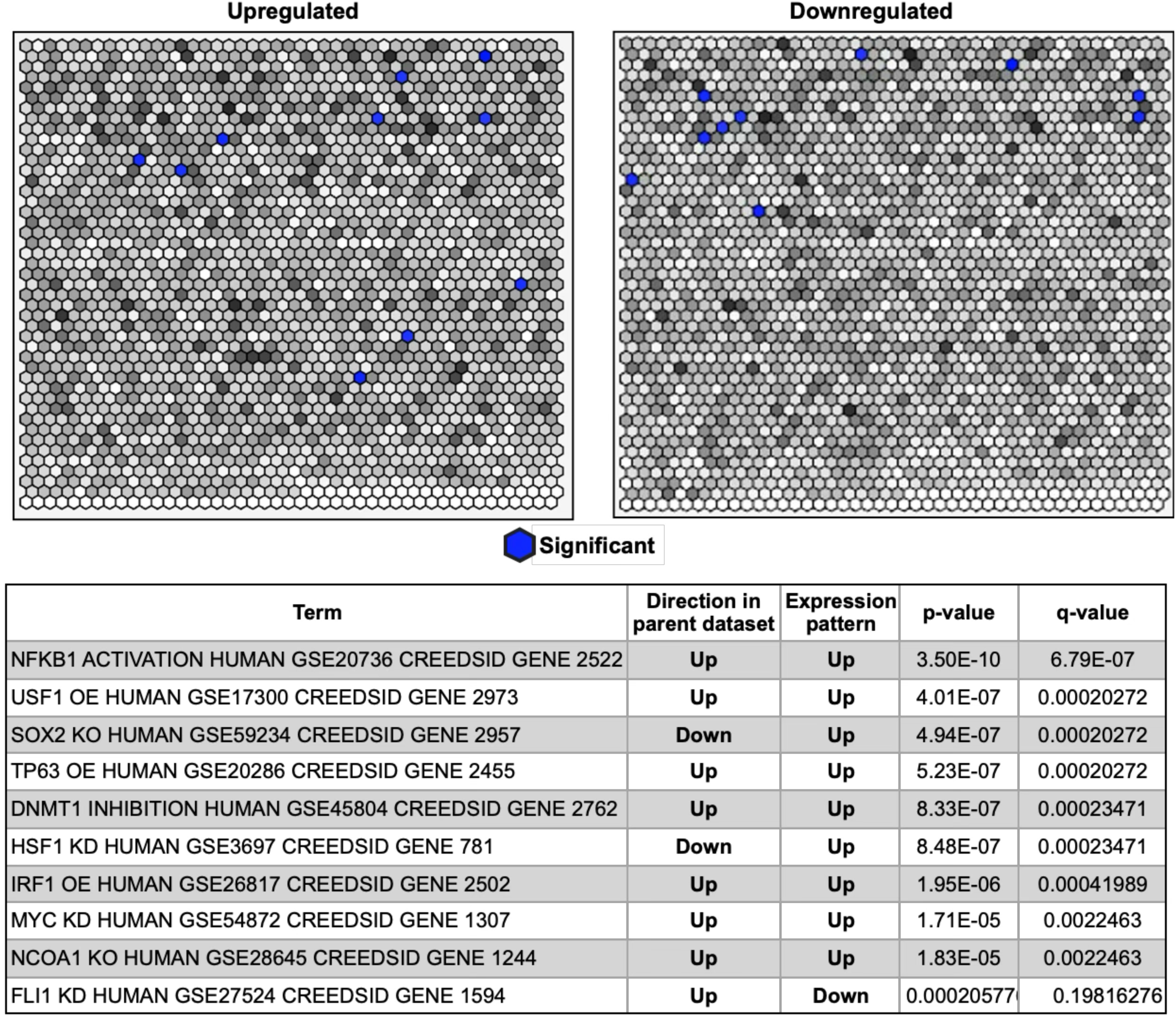
Hexagonal enrichment plot comparing gene set activation and silencing signatures in ATN1(DRPLA) versus ATN1(WT) (A) Gene set enrichment analysis was performed using ENRICHR to identify transcriptional similarity between differentially expressed genes in the ATN1(DRPLA) versus ATN1(WT) comparison and curated perturbation-based gene signatures. Each hexagon in the plot represents a single term. The brighter the color, the higher the Jaccard similarity between the term gene set and the input gene set. Similar gene sets are generally grouped together. The terms highlighted in blue are the most significantly overlapping with the input query gene set. (B) Highlighted gene sets include transcriptional programs linked to NFKB1, IRF1, USF1, TP63, SOX2, HSF1, and FLI1, revealing convergence of ATN1(DRPLA)-induced expression changes with known inflammatory, chromatin remodeling, stress-response, and differentiation pathways.

Gene sets typically downregulated following SOX2 and HSF1 knockdown were instead upregulated in our dataset, implicating activation of these key regulators and suggesting a compensatory induction of differentiation and stress-response programs in ATN1(DRPLA) cells. Conversely, transcriptional profiles associated with FLI1 knockdown were downregulated in our dataset, pointing to a distinct regulatory trajectory. Collectively, these findings highlight the broad reprogramming of transcriptional networks in DRPLA and underscore the involvement of diverse regulatory axes in disease pathogenesis.

### Proteotoxic stress and chaperone dysregulation in DRPLA

Given the evidence of HSF1 activation (figure 5) and its well-established role in regulating protein quality control pathways including those disrupted in other polyQ diseases (Kampinga & Bergink, 2016; Singh et al., 2024; Wyttenbach, 2004), we assessed whether heat shock proteins, small heat shock proteins, and protein chaperones were differentially expressed in ATN1(DRPLA) compared to ATN1(WT). In total, 34 heat shock- and chaperone-related genes were upregulated, while 21 were downregulated, reflecting a complex regulatory landscape (figure 6A). Overall, chaperones involved in endoplasmic reticulum-associated degradation (ERAD) pathway were downregulated and markedly absent from the upregulated chaperone cohorts. The predominance of upregulated chaperone genes supports the presence of proteotoxic stress in the ATN1(DRPLA) model. We also observed decreased expression of DNAJB12/DNAJB14 which promotes maturation of potassium channels including KCND2, amongst other functions. Furthermore, we observed downregulation of DNAJB6, a critical molecular chaperone involved in maintaining cytoskeletal integrity through organization of KRT8/KRT18 filaments. DNAJB6 acts as a molecular chaperone for neuronal proteins, including HTT (Smith & D’Mello, 2016).

**Figure 6.**
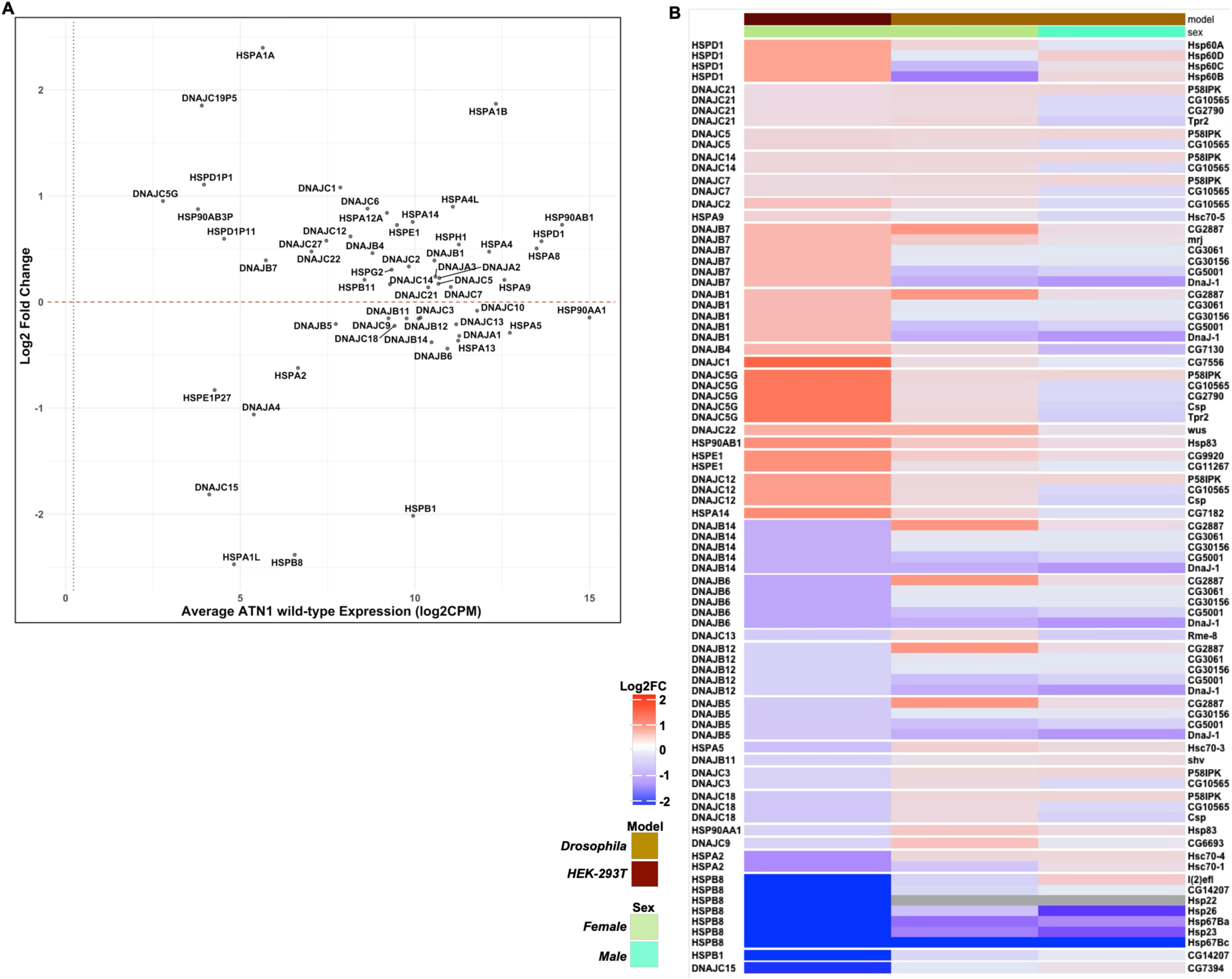
Chaperone expression changes in ATN1(DRPLA) versus ATN1(WT) and their conservation across species. (A) Bivariate plot displaying transcriptional changes in heat shock proteins and molecular chaperones. The X-axis shows the average normalized expression (trimmed mean TPM) of each gene in the ATN1(WT) condition, while the Y-axis represents the log2 fold change in ATN1(DRPLA) versus ATN1(WT). Genes above the horizontal line are transcriptionally upregulated in the DRPLA condition, while those below are downregulated. (B) Heatmap of HSPs and chaperones differentially expressed in ATN1(DRPLA) versus ATN1(WT), annotated with orthologs from Homo sapiens (left annotation column) and *Drosophila melanogaster* (right annotation column), as determined using BioMart. For human genes with multiple *Drosophila* orthologs, each ortholog is shown individually but grouped on a single row to reflect shared ancestry.

Previous work from our group characterized the molecular landscape of wild-type ATN1 and polyQ-expanded ATN1 (Q88) in *Drosophila melanogaster*, revealing sex-specific dysregulation of protein quality control pathways (Prifti et al., 2025). Building on these findings, we compared differentially expressed protein quality control genes from our HEK293T ATN1(DRPLA) cells to their corresponding orthologs in the *Drosophila* model. In several cases, the expression patterns of heat shock protein orthologs were conserved across models, with strong similarities between HEK293T cells, which are female in origin, and female flies (figure 6B). These findings indicate that polyQ-expanded ATN1 induces proteotoxic stress and disrupts protein quality control networks, with conserved chaperone expression changes observed across human cell and *Drosophila* models.

### Enhanced oxidative stress sensitivity in ATN1(DRPLA) cells

Oxidative stress has been implicated in DRPLA pathology based on analyses of patient tissue (Miyata et al., 2008). To investigate whether polyQ-expanded ATN1 alters redox homeostasis in our cell model, we examined the expression of genes involved in reactive oxygen species (ROS) metabolism. Differential expression analysis revealed a mixed oxidative stress signature in ATN1(DRPLA) cells compared to ATN1(WT). We observed upregulation of PFKP, a key glycolytic enzyme linked to ROS generation through metabolic reprogramming; MGST1, a microsomal glutathione S-transferase associated with detoxification of lipid peroxides; and MBP, myelin basic protein, which has been implicated in oxidative injury pathways. In contrast, NQO1, an NAD(P)H dehydrogenase with antioxidant function, and IPCEF1, involved in cytoskeletal responses to oxidative stress, were significantly downregulated (figure 7A), suggesting impaired antioxidant capacity.

**Figure 7.**
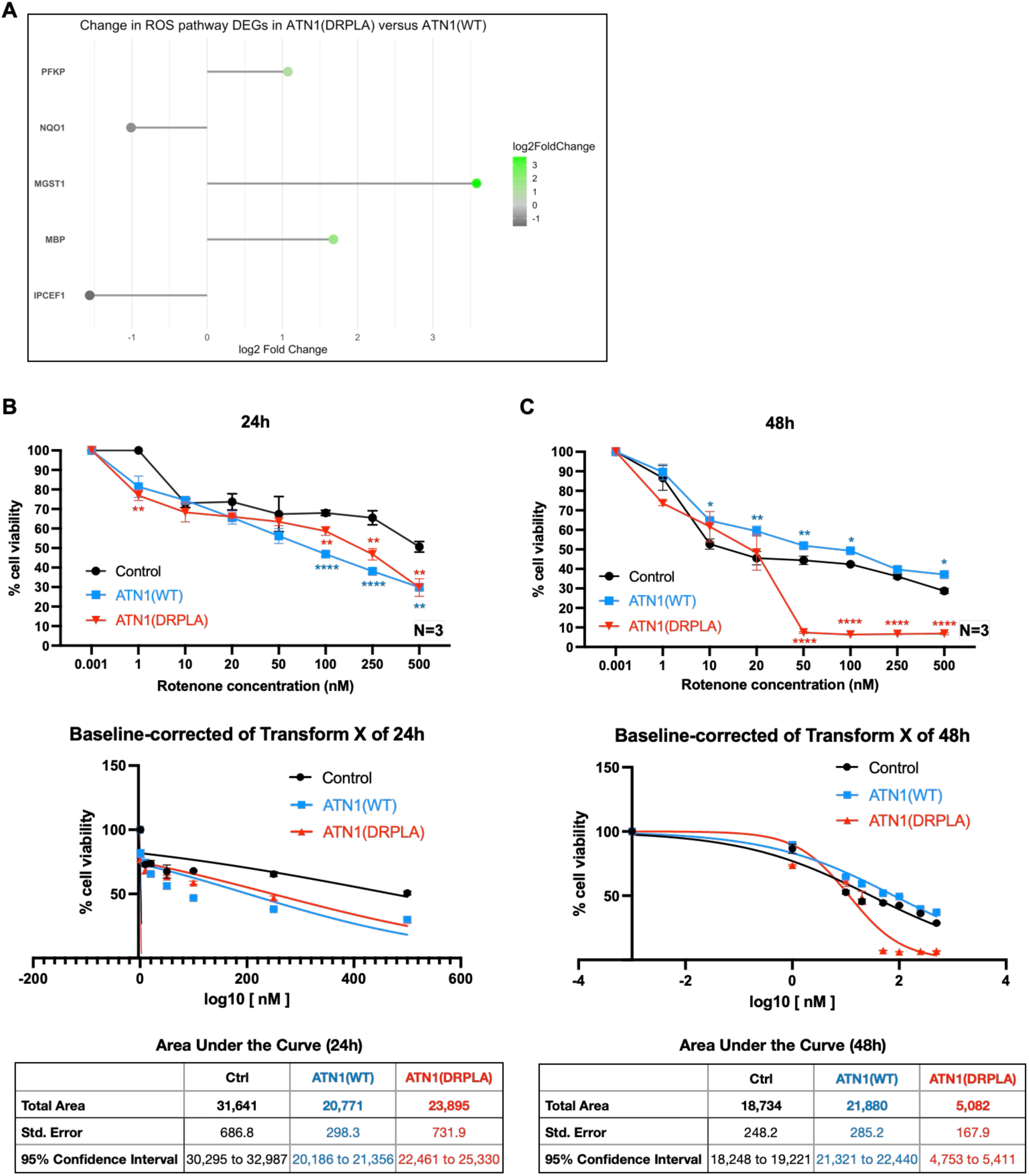
Increased susceptibility to mitochondrial oxidative stress in ATN1(DRPLA) cells. (A) Lollipop plot displaying log2 fold changes (x-axis) of reactive oxygen species (ROS)-related genes differentially expressed between ATN1(DRPLA) and ATN1(WT). Black dots denote genes that are downregulated in ATN1(DRPLA), while green dots indicate genes that are upregulated. (B) and (C), MTT cell viability assays when HEK293T cells stably expressing the indicated constructs were exposed to increasing concentrations of rotenone for 24 (B) or 48 (C) hours. Data are graphed as both viability (top portions) and baseline-corrected transformations of X. Asterisks in panels are from two-way ANOVAs, where *, P<0.05, **, P<0.01, ***, P<0.001, ****, P<0.0001. N=3 independent repeats.

This imbalance between ROS-generating and ROS-detoxifying genes pointed to a potential redox vulnerability in the DRPLA model. To probe mitochondrial sensitivity to oxidative stress, we treated cells with rotenone, a complex I inhibitor known to elevate mitochondrial ROS levels. We treated control, ATN1(WT), and ATN1(DRPLA) stable cell lines with increasing concentrations of rotenone (0 to 500 nM) for 24 and 48 hours (figure 7B, C). Cell viability was measured using the MTT assay and normalized to each group’s untreated condition. At low rotenone concentrations (≤10 nM), all cell lines maintained viability. However, as concentrations increased (≥100 nM) after 24h, ATN1(WT) and ATN1(DRPLA) cells exhibited a significant decline in viability compared to the control group (figure 7B). After 48h, across several concentrations (starting at 50nM), ATN1(DRPLA)-expressing cells displayed significantly lower viability than both ATN1(WT) and control (p < 0.01) (figure 7C). The nonlinear dose–response curve showed that ATN1(DRPLA)-expressing cells exhibited a sharper decline in viability compared to ATN1(WT), particularly at 48 hours (figure 7B, C). These data suggest increased susceptibility of ATN1(DRPLA)-expressing cells to mitochondrial stress induced by rotenone.

## DISCUSSION

We developed HEK293T cell lines stably expressing wild-type or polyQ-expanded ATN1. Transcriptomic profiling revealed that ATN1(DRPLA) expression induces widespread gene expression changes affecting synaptic function, ion channel activity, and extracellular matrix remodeling. PolyQ expansion in ATN1 also disrupted transcriptional regulatory networks, activated proteotoxic stress responses, and impaired redox homeostasis in cells. Functional assays showed increased sensitivity of ATN1(DRPLA) cells to mitochondrial oxidative stress, and cross-species analysis revealed conserved chaperone expression patterns. Together, these findings highlight convergent pathogenic mechanisms and identify potential molecular targets for therapeutic intervention in DRPLA.

PolyQ-expanded ATN1 induces a distinct gene expression program in HEK293T cells, diverging from both control and wild-type ATN1 conditions. Modest increases in ATN1 expression, regardless of polyQ expansion, altered the expression of genes involved in chromatin remodeling, proteostasis, and cytoskeletal organization suggesting that transcriptional dosage alone may contribute to cellular vulnerability. The fact that several genes were similarly affected in both ATN1(WT) and ATN1(DRPLA) conditions further supports the notion that ATN1 overexpression imposes a basal level of transcriptional stress, which may sensitize cells to the additional toxicity conferred by the polyQ expansion.

Mild overexpression of ATN1(WT) appears to support neural-like gene expression, whereas polyQ-expanded ATN1 drives a shift toward aberrant remodeling, with disruption of synaptic signaling, altered calcium dynamics, and impaired ion transport—features characteristic of early pathogenic processes in polyQ disorders such as HD and SCAs (Chopra et al., 2020; Huang et al., 2025; Lieberman et al., 2019; Luttik et al., 2022; Moreira-Gomes & Nóbrega, 2024; Tejwani et al., 2024; H. Zhang & Wang, 2024). Indeed, our analysis suggests that ATN1(DRPLA) expression was associated with an elevated stem-like transcriptional signature along with increased expression of MYC (log2FC >1.5) (supplemental tables 1 and 2) and the enrichment of SOX2 gene activation signature (figure 5), whereas wild-type ATN1 appeared to promote a more differentiated profile (supplemental figure 5). This observation stands in contrast to the notion that neurodegeneration and brain aging are accompanied by a progressive loss of stem-like features and regenerative capacity in precursor cells (Encinas & Sierra, 2012; M. Li, Guo, Carey, & Huang, 2024; Ruetz et al., 2024). The inverse phenotype observed here may be explained by the absence of aggregated protein. Without direct functional assays, it remains unclear whether this state confers plasticity or instead represents a stalled or aberrant cellular identity. These findings suggests that polyQ-expanded ATN1 maintains or reactivates developmental transcriptional programs, potentially as a compensatory or stress-responsive mechanism. It also raises the possibility that polyQ expansions in ATN1 affect not only transcriptional repression activity but also the broader cellular state, reshaping how differentiation cues are integrated or implemented.

Further enrichment analyses revealed transcriptional activation of inflammation- and stress-related programs, including targets of NFKB1, IRF1, and HSF1, and gene sets typically downregulated upon SOX2 and HSF1 knockdown, suggesting compensatory activation of differentiation and proteostasis networks. In contrast, the downregulation of FLI1-responsive genes points to a divergent regulatory axis uniquely affected in the DRPLA context. Consistent with these findings, we observed widespread upregulation of heat shock proteins and molecular chaperones in the ATN1(DRPLA) model, indicative of proteotoxic stress. However, key components of the ER-associated degradation (ERAD) pathway and cytoskeletal chaperones such as DNAJB6 and DNAJB12 were downregulated, suggesting selective impairment of critical proteostasis mechanisms. Expression patterns of several chaperone genes were conserved between our human cell model and female *Drosophila* expressing expanded ATN1, highlighting biologically relevant and potentially sex-specific aspects of the stress response to polyQ expansion. Moreover, chromosomal enrichment analysis of ATN1(DRPLA) *versus* ATN1(WT) DEGs were enriched for X-linked genes (supplemental figure 6). The enrichment of X-linked genes among DEGs in ATN1(DRPLA) *versus* ATN1(WT) suggests that polyQ-expanded ATN1 disproportionately impacts regulatory elements on the X chromosome. Given the X chromosome’s unique dosage regulation and enrichment for neurodevelopmental genes (Nguyen & Disteche, 2006; Yin et al., 2009; Zechner & Hameister, 2011), this finding raises the possibility that DRPLA pathogenesis involves disrupted control of sex-linked transcriptional programs. Such dysregulation could contribute to sex-biased symptom severity or even onset, particularly among individuals with comparable CAG repeat lengths, and may underlie aspects of developmental derailment or circuit-level dysfunction. These findings underscore the need to further investigate chromosomal context especially X-linked regulatory mechanisms in ATN1(DRPLA) disease models.

In HEK293T cells expressing ATN1(DRPLA), we observed transcriptional silencing of MLH3, a key component of the DNA mismatch repair (MMR) machinery. MLH3 forms the MutLγ complex with MLH1 and operates downstream of the MutSβ complex (composed of MSH2 and MSH3) to resolve insertion/deletion loops and trinucleotide repeat mismatches (Yin et al., 2009) (Benn, Gibson, & Reynolds, 2021; A. Shibata & Jeggo, 2020). In contrast, increased MSH3 expression has been implicated in genome-wide association studies (GWAS) of HD as a modifier of somatic CAG repeat instability, and its suppression has yielded protective outcomes in HD models (Bunting et al., 2025; Flower et al., 2019; Moss et al., 2017). These findings suggest that MMR components like MSH3 and MLH3 play divergent yet complementary roles in modulating trinucleotide repeat stability. The silencing of MLH3 in our DRPLA model may reflect an adaptive suppression of error-prone repeat repair, potentially limiting further expansion but at the cost of reduced DNA repair fidelity. This is notable given our concurrent observation of activation of DNA damage marker RPRM, a p53-regulated gene associated with G2/M arrest and cellular stress responses. These data point to disease-specific alterations in the DNA repair landscape in DRPLA and highlight the need to further dissect the functional interplay between MMR components in polyglutamine expansion disorders.

Our expression overlay network (supplemental figure 7) highlights several transcriptionally activated genes in ATN1(DRPLA) that are also pharmacologically targetable, providing a foundation for potential therapeutic intervention. Notably, PDGFD, a growth factor upregulated in the HEK293T cells expressing ATN1(DRPLA), signals through PDGFR-β and may be indirectly inhibited using receptor tyrosine kinase (RTK) inhibitors such as imatinib or crenolanib (Iqbal & Iqbal, 2014; Tinkle et al., 2021; P. Wang et al., 2014). Similarly, GRIK3, a kainate-type glutamate receptor subunit implicated in ASD, could be modulated with selective kainate receptor antagonists (Jane, Lodge, & Collingridge, 2009; Motazacker et al., 2007; H. Shibata et al., 2006). We additionally identified IFIH1 (MDA5) as a prominent immune-related node, which signals via the JAK-STAT axis and could be targeted using JAK inhibitors (e.g., ruxolitinib) (Maiti, 2024; Rodero & Crow, 2016; Schwartz et al., 2017a, 2017b). Inflammatory transcription factors like CEBPD may be indirectly suppressed by targeting upstream regulators such as STAT3 or NF-κB. These pharmacologically relevant nodes converge on key pathological processes including inflammation, aberrant excitatory signaling, and proliferative signaling cascades, all of which have been implicated in neurodegeneration.

Prior research utilizing transient expression of ATN1 in COS-7 and Neuro2a cells found that a mutant C-terminal fragment of ATN1 containing an expanded polyQ tract selectively accumulated in cells. This fragment localized to cytoplasmic membranes, organelles, and insoluble fractions. Its accumulation, influenced by caspase activity, suggested that proteolytic processing of ATN1 drives pathological protein aggregation and cellular dysfunction in DRPLA (Suzuki et al., 2010). Transient expression of full-length ATN1 in differentiated PC12 cells through adenoviruses led to intranuclear inclusions, although their presence did not correlate with increased toxicity (A. Sato et al., 1999). Mouse- and fly-based studies from others and us were informative in helping to understand properties of ATN1 protein localization, aggregation, cleavage and function; its effects on electrophysiological anomalies; and the importance of CAG repeat length in disease severity (Napoletano et al., 2011; Prifti et al., 2025; Sakai et al., 2006; Sato et al., 2008; T. Sato et al., 1999; Schilling et al., 1999; Ying et al., 2006). Earlier work also identified perturbations in transcription, synaptic transmission, and protein quality control related to pathogenic ATN1, similar to our observations in this work, indicating important complementarity among the different models of DRPLA. The depth of the information that we presented here may now be used to further inform understanding of normal ATN1 function and DRPLA biology of disease, especially when considering changes in transcription profiles, protein quality control, and, particularly intriguingly, changes in “stemness” index.

We confirmed a handful of our transcriptomics results with WBs. While there was some data congruence, we also observed instances of incongruity. These findings underscore the need of future studies using proteomic and microscopy research to overlay transcriptomics, proteomics, and localization landscapes — especially in cases of proteins that are heavily modulated through posttranslational modifications (like ATXN3 in this report) — to pinpoint the relation of gene expression changes to the levels, localization, and activity of the proteins they encode.

We were intrigued by the data that ATN1, whether wild-type or pathogenic, led to changes in the expression of most polyQ disease protein genes. These effects at the transcript level did not consistently correlate to the same degree of change or direction in protein levels. But, they provided new clues into the possible role of ATN1 more globally in polyQ diseases, and warrant future investigation to understand how both normal and disease-causing ATN1 may be involved in the normal biology of polyQ disease genes and in their pathology.

We selected HEK293 cells due to their favorable properties for manipulation and protein expression analysis. Their high transfection efficiency and compatibility with a broad range of molecular tools allow for efficient delivery and sustained expression of constructs, facilitating different assays. The cells proliferate efficiently under standard culture conditions, are highly compatible with different assays, and support stable line generation making them well suited for reproducible experiments. HEK293 cells have a long-standing track record in biomedical research, supported by extensive protocols and validated reagents (Pulix et al., 2021; Stepanenko & Dmitrenko, 2015; Tan et al., 2021; Thomas & Smart, 2005). These characteristics make the HEK293 family of cells a practical model for the initial investigation of gene and protein function, including screening approaches to evaluate the cellular consequences of pathogenic ATN1 overexpression. However, the limitations of using HEK293 cells as a disease model must also be acknowledged. As a non-neuronal, immortalized cell line, HEK293T cells lack the specialized transcriptional environments found in neurons. This may limit the relevance of observed phenotypes to those that occur in DRPLA-affected tissues. Moreover, exogenous expression systems often drive ATN1 production at levels that exceed physiological norms, potentially disrupting native regulatory mechanisms and obscuring dosage-sensitive cellular responses. HEK293 cells may also differ in their handling of misfolded proteins, with altered aggregation kinetics and cytotoxic responses compared to differentiated neuronal cells. While these constraints should be considered when interpreting results, the use of HEK293 cells enabled precise manipulation of ATN1 expression and facilitated hypothesis generation for validation in neuronally relevant models. The development of DRPLA patient-derived iPSCs may be of particular relevance in this direction (Bidollari et al., 2019; Kozlowska, Ciolak, Olejniczak, & Fiszer, 2019).

Future studies are clearly needed to further investigate our findings and place them in the context of DRPLA disease biology. Still, our current findings are helpful in understanding not only the impact of both normal and pathogenic ATN1 protein in the mammalian cellular environment, but also and in pinpointing specific leads for pursuit in these and other models of this incurable disease.

## ACKNOWLEDGEMENTS

We thank Mr. Ryan Dulay and Mr. Truman Kula for their assistance with Western blotting. This study was funded in part by a Pathway to Faculty award to ON from the Provost of Wayne State University, by a postdoctoral award to MP from the Office of the Vice President of Research at Wayne State University, from by R01NS086778-10A1S1 to SVT from NINDS, and by R01NS086778 to SVT from NINDS.

## AUTHORS’ CONTRIBUTIONS

***ON:*** conceptualization, data curation, software, formal analysis, funding acquisition, validation, investigation, visualization, methodology and writing and editing.

***MP:*** conceptualization, data curation, software, formal analysis, funding acquisition, validation, investigation, visualization, methodology and writing and editing.

***JPR:*** data curation, software, formal analysis, validation, investigation, visualization, methodology and writing and editing.

***SVT:*** conceptualization, resources, data curation, software, formal analysis, supervision, funding acquisition, validation, investigation, visualization, methodology and writing and editing.

All authors contributed to the article and approved the submitted version.

