## Supplemental figure 1 for "Disrupted Transcriptional Networks by Mutant Atrophin-1 in a Cell Culture Model of Dentatorubral-Pallidoluysian Atrophy"

### SUPPLEMENTAL FIGURE LEGENDS

**Supplemental figure 1: RNA-seq quality control metrics for individual samples.**

Mean quality scores for each base position across all paired-end reads are shown for each RNA-seq sample. Quality scores were assessed using FastQC and plots represent the per-base sequence quality distribution for both forward and reverse reads.

**Supplemental figure 2: Soluble/pellet fractionation using NETN lysis buffer.**

A) Fluorescence images similar to figure 1F.

B) Outcomes from soluble/pellet preparation of cells expressing the indicated constructs. Shown in graph are means -/+ SD. Statistics: not significant (ns) by Brown-Forsythe and Welch ANOVA tests. To calculate each data point in the graph, ATN1 signal was divided by its respective Direct Blue lane signal, then normalized to the fraction of total lysate (soluble) or resuspension (pellet) fraction that it represents, and expressed as a fraction of 'soluble/(soluble+pellet) signal' for that specific independent repeat.

**Supplemental figure 3: Expression of polyQ disease protein-encoding genes across experimental conditions.**

Box plots with variable axis scaling showing the mRNA expression levels (TPM) of selected polyQ-related genes across all conditions: control, ATN1(WT), and ATN1(DRPLA). Each box represents data from N=5 independent replicates per condition. Note: CACNA1A expression levels were below the typical threshold for low-count filtering and would ordinarily be excluded from differential expression analysis due to limited read depth. However, these data are retained for completeness in representing polyQ disease-associated genes. We advise caution in interpreting its expression values.

**Supplemental figure 4: WB validation of protein expression across experimental conditions.**

Representative WB and quantification of target protein levels in HEK-293T cells expressing control, ATN1(WT), or ATN1(DRPLA). Protein lysates from N=5 biological replicates per condition were analyzed, except ‘Ataxin2’ which was N=9. Bands were normalized to corresponding total protein loading (Direct Blue). Shown are mean -/+ SD. Statistics, Brown-Forsythe and Welch ANOVA, expect Ataxin-2, which was assessed by Kruskal-Wallis. ns, not significant, *, P<0.05, **, P<0.01.

**Supplemental figure 5: Stemness index (mRNAsi) calculated for each condition.**

Box plot showing the distribution of mRNAsi scores for each condition: control, ATN1(WT), and ATN1(DRPLA). Scores were calculated using a one-class machine learning model trained on stem cell expression profiles, as described by Malta et al. (2018). Each point represents an individual biological replicate (N=5). Boxes represent the interquartile range (IQR) and lines indicating the median.

Malta TM, Sokolov A, Gentles AJ, Burzykowski T, Poisson L, Weinstein JN, Kamińska B, Huelsken J, Omberg L, Gevaert O, Colaprico A, Czerwińska P, Mazurek S, Mishra L, Heyn H, Krasnitz A, Godwin AK, Lazar AJ; Cancer Genome Atlas Research Network; Stuart JM, Hoadley KA, Laird PW, Noushmehr H, Wiznerowicz M. Machine Learning Identifies Stemness Features Associated with Oncogenic Dedifferentiation. Cell. 2018 Apr 5;173(2):338-354.e15. doi: 10.1016/j.cell.2018.03.034. PMID: 29625051; PMCID: PMC5902191.

**Supplemental figure 6. Chromosomal enrichment analysis of DEGs between ATN1(DRPLA) and ATN1(WT).**

Forest plot showing odds ratios (OR) and 95% confidence intervals (CI) for enrichment of differentially expressed genes (DEGs) on each human chromosome in the ATN1(DRPLA) *versus* ATN1(WT) comparison. Enrichment was assessed using Fisher's exact test, comparing the observed versus expected number of DEGs per chromosome. Points represent the OR for each chromosome relative to genomic expectation, with horizontal lines indicating the 95% CI. Mustard-colored points indicate chromosomes significantly enriched for DEGs (OR>1, P<0.05), blue points indicate significantly depleted chromosomes (OR<1, P<0.05), and black points represent chromosomes with no significant enrichment.

**Supplemental figure 7: Targetable Gene Network in genes 'turned-on' in ATN1(DRPLA) relative to ATN1(WT).**

The network illustrates pharmacologically targetable genes identified from the ATN1(DRPLA) *versus* ATN1(WT) differential expression dataset, overlaid with $\log_2$ fold-change values. Italic text below nodes: known small molecule inhibitors or modulators.

### Supplemental figure 1

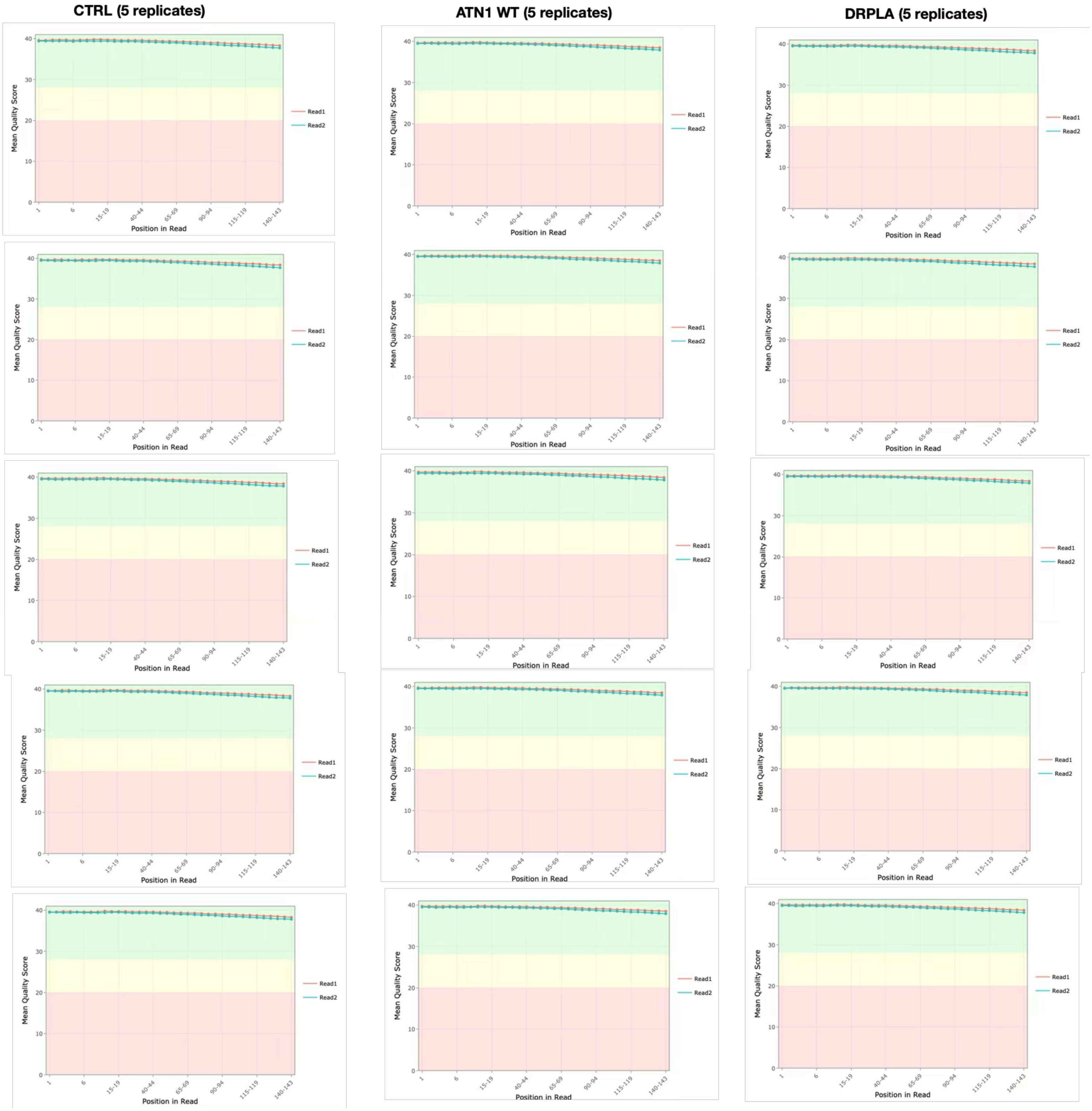

### Supplemental figure 2

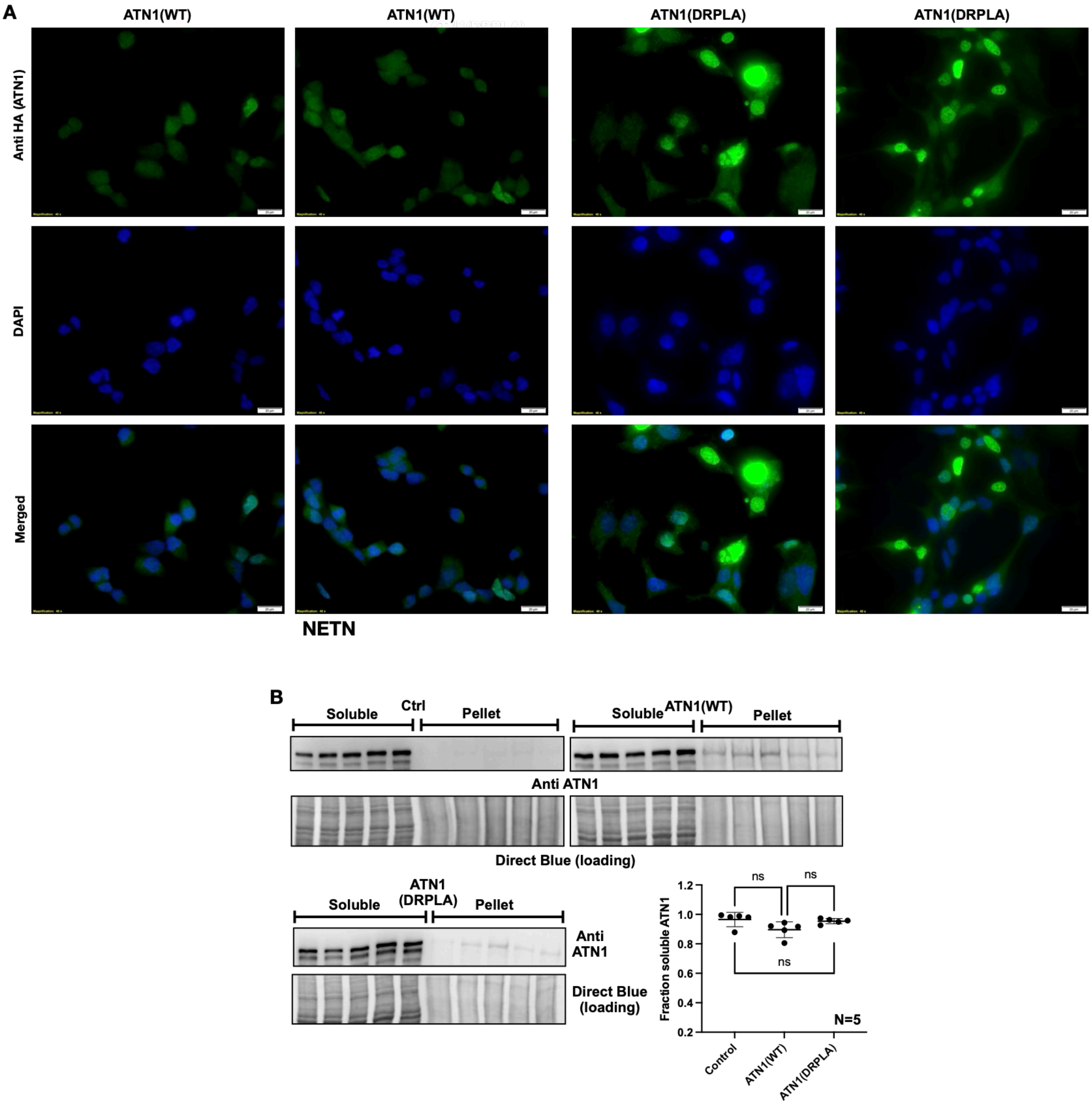

**Supplemental figure 3**

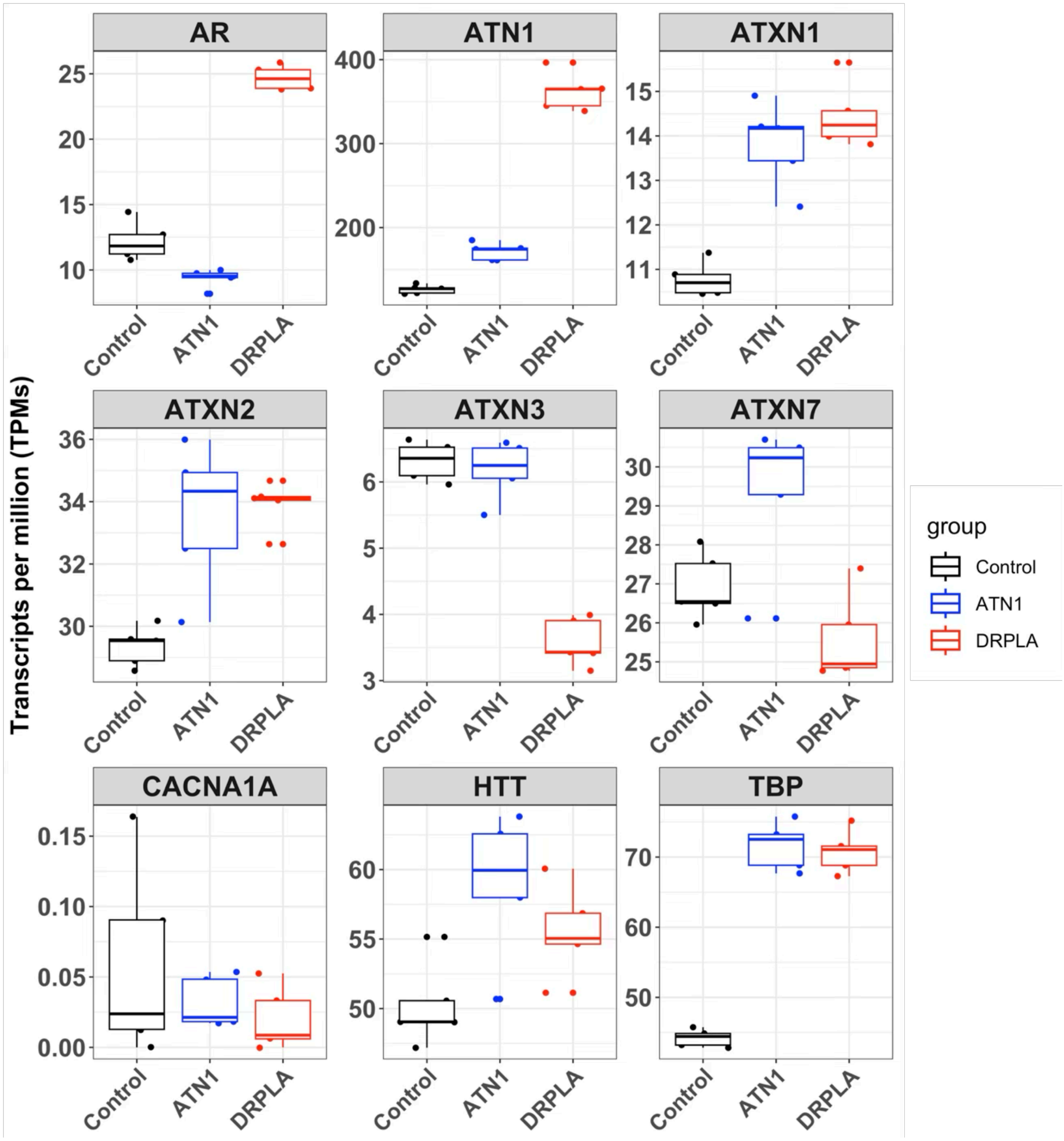

**Supplemental figure 4**

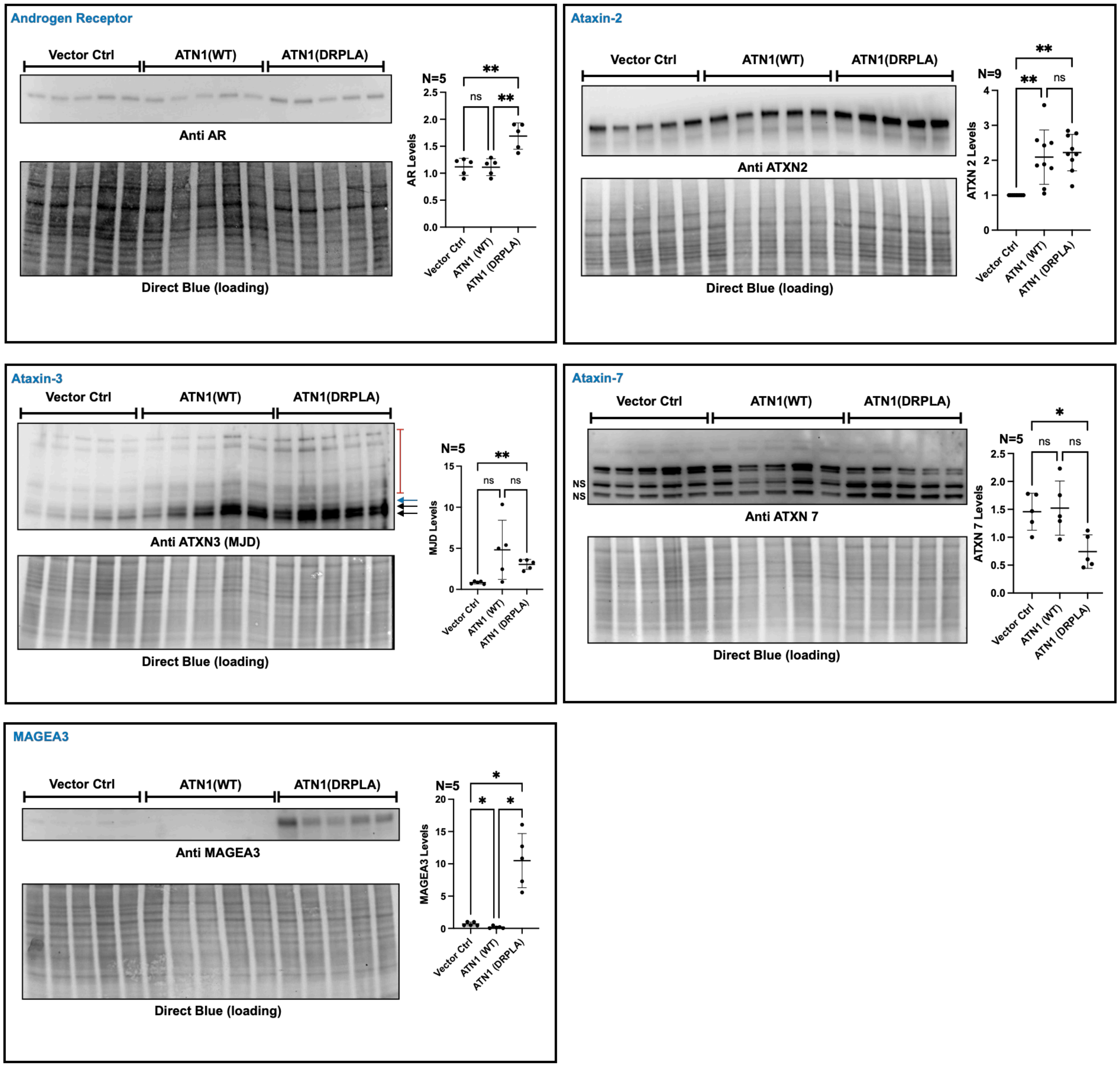

**Supplemental figure 5**

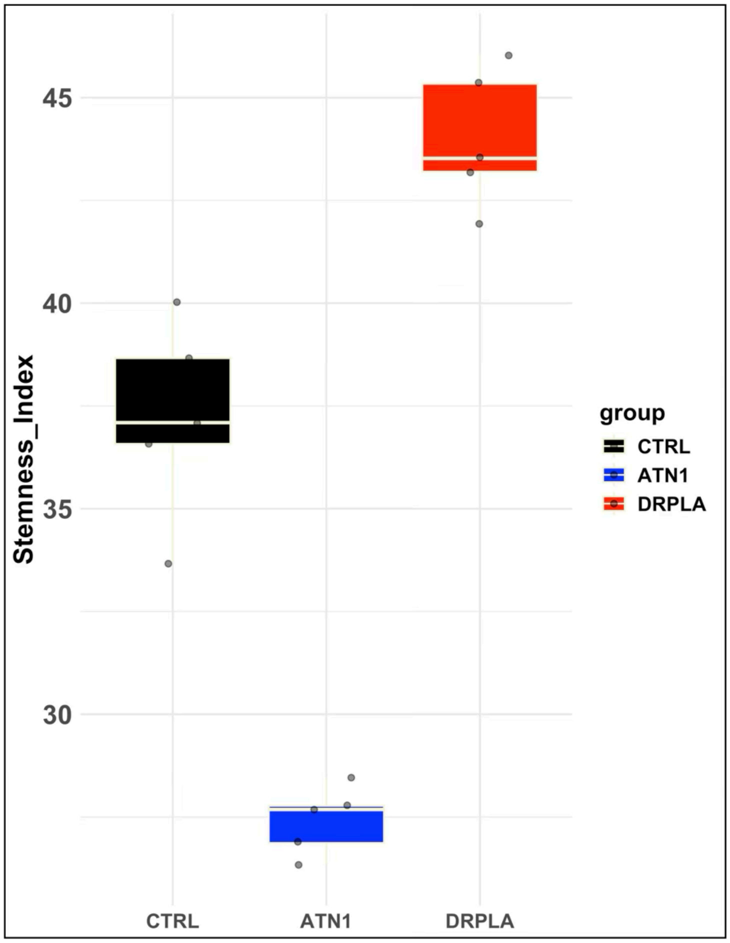

**Supplemental figure 6**

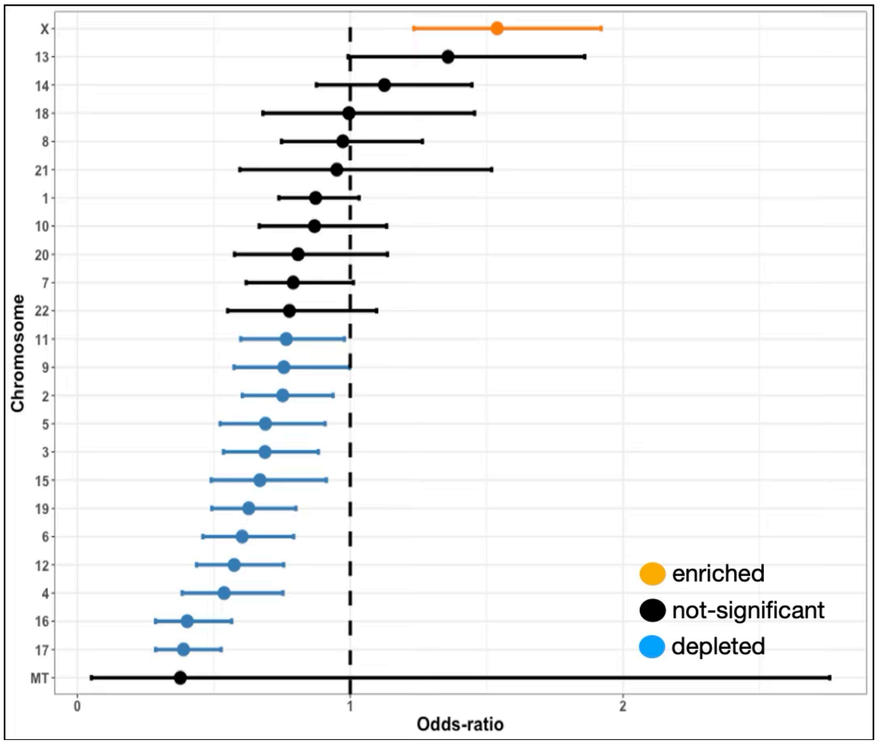

### Supplemental figure 7

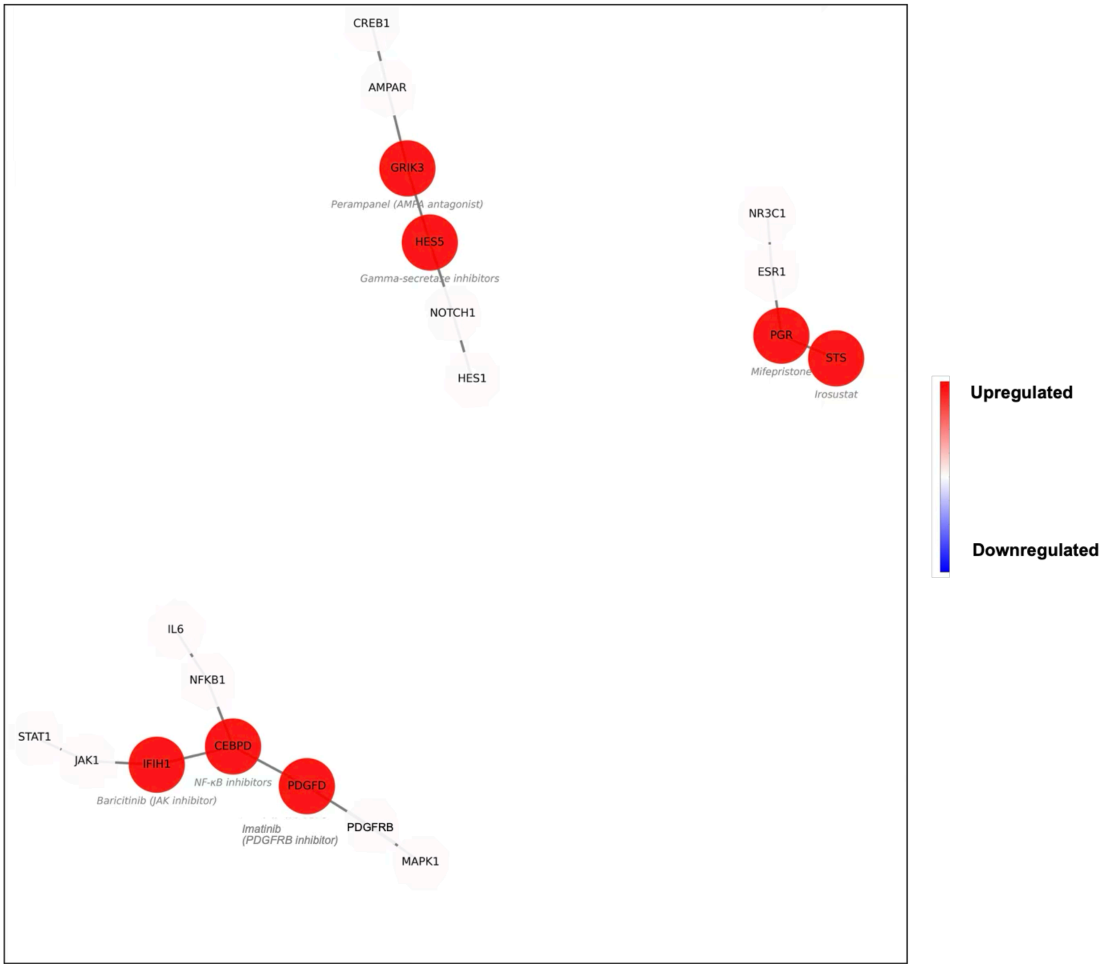
